# Piezo1-mediated spontaneous activity of satellite glia impacts DRG development

**DOI:** 10.1101/2022.09.28.509897

**Authors:** Jacob P. Brandt, Cody J. Smith

## Abstract

Spontaneous activity of neural cells is a hallmark of the developing nervous system. It is widely accepted that chemical signals, like neurotransmitters, contribute to spontaneous activity in the nervous system. Here we reveal an additional mechanism of spontaneous activity that is mechanosensitive in the peripheral nervous system (PNS) using intravital imaging of growing dorsal root ganglia (DRG) in zebrafish embryos. GCaMP6s imaging shows that developing DRG satellite glia contain distinct spontaneous Ca^2+^ transients, classified into simultaneous, isolated and microdomains. Longitudinal analysis over days in development demonstrates that as DRG satellite glia become more synchronized, isolated Ca^2+^ transients remain constant. Using a chemical screen, we identify that Ca^2+^ transients in DRG glia are dependent on mechanical properties, which we confirmed using an experimental application of mechanical force. We find that isolated spontaneous Ca^2+^ transients of the glia during development is altered by manipulation of mechanosensitive protein Piezo1, which is expressed in the developing ganglia. In contrast, simultaneous Ca^2+^ transients of DRG satellite glia is not Piezo1-mediated, thus demonstrating that distinct mechanisms mediate subtypes of spontaneous Ca^2+^ transients. Activating Piezo1 eventually impacts the cell abundance of DRG cells and behaviors that are driven by DRG neurons. Together, our results reveal mechanistically distinct subtypes of Ca^2+^ transients in satellite glia and introduce mechanobiology as a critical component of spontaneous activity in the developing peripheral nervous system.

## INTRODUCTION

It is widely accepted that spontaneous activity is a critical feature of the developing nervous system[1–3]. This activity has been visualized by measuring Ca^2+^ transients. For years, such spontaneous Ca^2+^ transients have been investigated in neurons and are identified as neuronal firing or activity, but recent studies have also revealed an important role for spontaneous Ca^2+^ transients in glia. These glial Ca^2+^ transients can be in response to neuronal activity or independent of neuronal activity and can be characterized into distinct subtypes[4–7]. For example, glial cells exhibit whole cell and microdomain Ca^2+^ transients, which are mechanistically and functionally distinct[8–10]. Glial cells can also exhibit synchronous Ca^2+^ transients in physically-connected networks[11,12]. Regardless of Ca^2+^ transient subtype and unique from neurons, immature glia and their progenitors also proliferate throughout life[13]. How glial Ca^2+^ transients, proliferation and physically-connected networks are related or regulated, remains largely unexplored. The importance of these concepts is underscored by the prevalence of such processes during normal brain development and in gliomas[4,5,14–16]. If glial Ca^2+^ transients are critical for nervous system function, we need more investigation into how distinct Ca^2+^ transients change over development, whether different molecular components control distinct transient subtypes, and if distinct Ca^2+^ transients are linked to proliferation and/or network formation. Lastly, these concepts need to be explored in both the CNS and PNS.

What we do know is that spontaneous Ca^2+^ transients in the nervous system have largely been characterized to be dependent on chemical signals. In neurons, Ca^2+^ spontaneous activity is promoted by neurotransmitters and their receptors[17,18]. Similarly, glutamate and NMDA drive spontaneous Ca^2+^ transients of glial cells like oligodendrocytes and astrocytes[19–22]. We also know chemical signals like ATP can induce purinergic receptors to drive Ca^2+^ changes in glia, akin to activity of the glia[23–25]. Each of these chemical signals causes changes to ion channels that drive spontaneous Ca^2+^ transients. However, in addition to ion channels that are induced by chemical signals, mechanosensitive ion channels are also present in the nervous system[26,27]. For example, Piezo proteins are mechanosensitive channels that are expressed in the nervous system[26,28,29]. These mechanosensitive channels are essential for evoking a subset of peripheral sensory neurons in response to mechanical stimulation[30,31]. Peripheral mechanosensitive glia are also present at the skin to ensure response to mechanical stimuli[32]. However, the role of mechanosensitive properties in the development of glia is less understood, especially in peripheral glia. This is despite knowledge that mechanical components can have profound effects on cell differentiation and tissue organization and that Trp channels, some of which are at least partially mechanosensitive, are important for Ca^2+^ transients in glia like astrocytes[4,8,29,33–36].

Here we use imaging of GCaMP6s in satellite glia of the DRG in zebrafish as a model to investigate the role of glial activity in the developing peripheral nervous system. The DRG is required for somatosensory stimuli in the PNS and contains somatosensory neurons and satellite glia that ensheath those neurons. We identify that satellite glia display at least three types (microdomain, isolated, and simultaneous) of spontaneous Ca^2+^ transients in early phases of development. By mapping the GCaMP6s events, we identify that the DRG transitioned to synchronized Ca^2+^ transients early in development, demonstrating the formation of glial networks within the first three days of DRG construction. In a pilot screen and follow-up experimental manipulations, we identify mechanosensitive ion channel Piezo1 as a modulator of the isolated Ca^2+^ transients of satellite glia in development and identify that these satellite glia are mechanosensitive. Perturbation of Piezo1 causes not only changes in isolated Ca^2+^ transients of DRG satellite glia but also in their expansion and function, demonstrating a potential consequence to altering isolated glial Ca^2+^ transients during development. Together, we introduce the role of mechanosensitive ion channels in the spontaneous Ca^2+^ transients of the developing peripheral nervous system.

## RESULTS

### DRG satellite glia exhibit distinct Ca^2+^ transients

To understand if DRG satellite glia display spontaneous Ca^2+^ transients, we first explored the Ca^2+^ transients of DRG cells in intact ganglia using intravital imaging in zebrafish. To do this we imaged transgenic animals expressing GCaMP6s in distinct DRG cell populations. It is known that the DRG has both a population of neurons and satellite glia. To image both of these populations, we used animals expressing *Tg(sox10:gal4+myl7); Tg(uas:GCaMP6s); Tg(neurod:tagRFP).* Cells expressing RFP were identified as neurons, while the glial population expressed GCaMP6s with the absence of RFP (Fig 1A). To further confirm that these *sox10^+^ neurod^−^* cells were satellite glia we investigated the morphology of these cells (Sup Fig 2A-B). Using these triple transgenics and identifying morphological features of DRG cells, we found that both neurons and glia that ensheathed those neurons were present in the DRG during the developmental window examined. We define satellite glia in this report as *sox10^+^ neurod^−^* cells located in the DRG with ensheathing phenotypes.[37,38] In addition to these transgenics, we also imaged neurons in the DRG using *Tg(neurod:gal4+myl7); Tg(uas:GCaMP6s)*, which uses regulatory sequences of *neurod* that are expressed in DRG neurons. This allowed for another approach in which we only investigate the neuronal population. We imaged 3dpf (days post fertilization) animals expressing GCaMP6s at a 15 second interval for 1 hour, which allowed us to capture a three-dimensional view of 3-4 DRG. To define a Ca^2+^ transient event in a cell, we calculated the z score of the integrated density of fluorescence of each individual cell during that 1-hour time period. Time points with a z score greater than 2.58 (represents 99% confidence interval) were considered Ca^2+^ transient event-containing time points. If a Ca^2+^ transient event lasted for consecutive time points it was still considered one Ca^2+^ transient event. Scoring of the z score of GCaMP6s integral density of fluorescence over the 1 hour period revealed that all DRG displayed cells with spontaneous Ca^2+^ transients, with remarkable activity in satellite glia (Fig 1B,C). Within individual DRG, satellite glia displayed on average 3.600±1.78 Ca^2+^ transient events in a 1-hour period (n=25 cells, 14 DRG, 5 animals) (Sup Fig 1A). We also measured 2.364±1.12 Ca^2+^ transient events in the neuronal population in a 1-hour period (n=11 cells, 9 DRG, 4 animals) (Sup Fig 1A), indicating that both neurons and glia display Ca^2+^ transients during DRG construction. Ca^2+^ transients can be quick in cells, so it is possible that a 15 sec imaging interval under-represented the number of Ca^2+^ transients. To address this, we imaged smaller z-stacks but with short time intervals of 5 secs. These results revealed that Ca^2+^ transients in *sox10^+^* cells lasted on average 2.21 timepoints using 5 sec imaging intervals and therefore were generally captured with 15 sec intervals (n=412 Ca^2+^ transient events, 97 cells, 20 DRG, 5 animals) (Sup Fig 1B). We therefore utilized 15 sec imaging intervals throughout this manuscript, allowing us to image z-stacks that covered the entire DRG at each timepoint. With the lack of knowledge about glial activity in the DRG, we further investigated the developmental, molecular, and functional features of Ca^2+^ transients in *sox10^+^* satellite glia.

**Figure 1:**
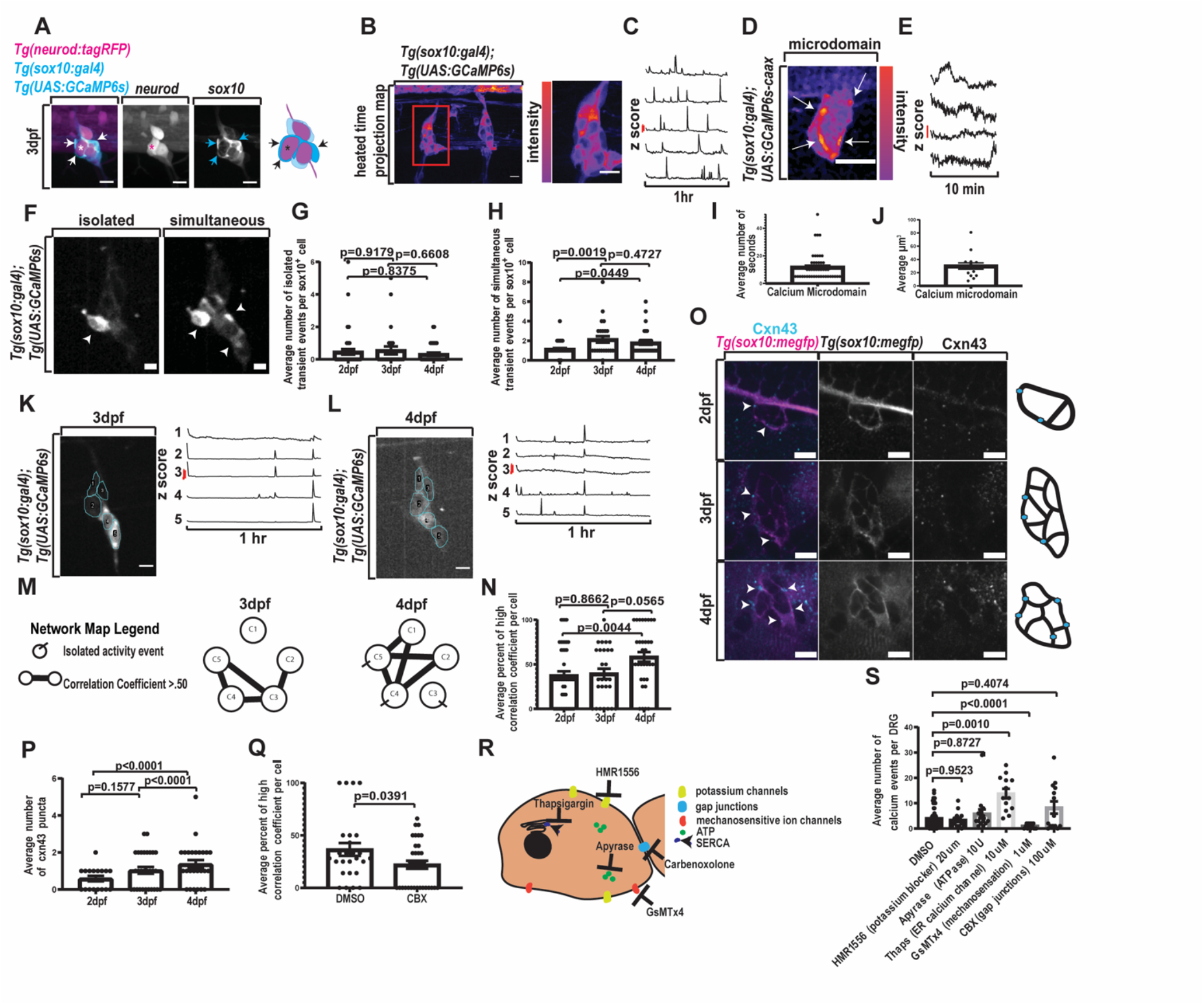
DRG exhibit distinct calcium activity and increase synchronous activity A) Confocal z-projections of DRG in a 3dpf animal expressing *Tg(sox10:gal4+myl7); Tg(uas:GCaMP6s); Tg(neurod:tagRFP).* Arrow notes satellite glia and asterisk notes neuron. B) Confocal z-projections of DRG in a 3 dpf animal expressing *Tg(sox10:gal4+myl7); Tg(uas:GCaMP6s)*. Images presented as heated scale (reds-more activity, blues-less activity). C) Line graphs depicting z scores of integrated density of fluorescence for individual cells expressing *Tg(sox10:gal4+myl7); Tg(uas:GCaMP6s)* over a 1 hour period. A z score greater than 2.58 indicates a Ca^2+^ transient event. Red scale bar is a z score of 2.58. D) Confocal z-projection of DRG in 3 dpf animal expressing *Tg(sox10:gal4+myl7); uas:GCaMP6s-caax*. Red colors indicate a higher intensity of fluorescence, and blue colors indicate a lower intensity of fluorescence. Arrows indicate active Ca^2+^ microdomains. E) Line graphs depicting z scores of integrated density of fluorescence for Ca^2+^ microdomains in 3dpf animals expressing *Tg(sox10:gal4+myl7); uas:GcaMP6s-caax* over a 10 minute period. A z score greater than 2.58 indicates an active Ca^2+^ event. Red scale bar is a z score of 2.58. F) Confocal z-projection of DRG in a 3 dpf animal expressing *Tg(sox10:gal4+myl7); Tg(uas:GcaMP6s)*. Left image depicts an isolated Ca^2+^ event, and the right image depicts a simultaneous Ca^2+^ event. Arrows indicate active cells. G) Average number of isolated Ca^2+^ transient events per *sox10^+^* cell at 2, 3, and 4 dpf in animals expressing *Tg(sox10:gal4+myl7); Tg(uas:GcaMP6s)*. (2dpf: 6 animals, 10 DRG, 46 cells, 3dpf: 4 animals, 6 DRG, 27 cells, 4dpf: 4 animals, 7 DRG, 34 cells) H) Average number of simultaneous Ca^2+^ transient events per *sox10^+^* cell at 2, 3, and 4 dpf in animals expressing *Tg(sox10:gal4+myl7); Tg(uas:GcaMP6s)*. (2dpf: 6 animals, 10 DRG, 46 cells, 3dpf: 4 animals, 6 DRG, 27 cells, 4dpf: 4 animals, 7 DRG, 34 cells) I) Average number of seconds for Ca^2+^ microdomains duration in 3dpf animals. (7 animals, 8 DRG, 15 microdomains) J) Average volume (μm^3^) of Ca^2+^ microdomains in 3dpf animals. (7 animals, 8 DRG, 15 microdomains) K-M) 3 (K) and 4 (L) dpf DRG in an animal expressing *Tg(sox10:gal4+myl7); Tg(uas:GCaMP6s)*. ROIs are traced for individual cells. Line graphs of the z score of the integrated density of fluorescence correspond to the individual ROIs. M) A corresponding network map for 3 and 4dpf DRG (K-L) indicates both the number of isolated Ca^2+^ transient events and the high correlation coefficients present in the DRG. N) Percent of high correlation coefficient per *sox10^+^* cell in animals expressing *Tg(sox10:gal4+myl7); Tg(uas:GCaMP6s)* at 2, 3, 4 dpf. (2dpf: 6 animals, 10 DRG, 50 cells 3dpf: 4 animals, 6 DRG, 27 cells 4dpf: 4 animals, 7 DRG, 34 cells) O) Immunohistochemistry for Cxn43 in DRG of animals expressing *Tg(sox10:meGFP)* at 2, 3, 4 dpf. Magenta indicates *Tg(sox10:meGFP)* and cyan indicates Cxn43. P) Quantification of the average number of Cxn43 puncta present in DRG at 2, 3, and 4 dpf. (2dpf: 6 animals, 18 DRG 3dpf: 10 animals, 29 DRG 4dpf: 8 animals, 24 DRG) Q) Quantification of the average percent of high correlation coefficients per *sox10^+^* cell following treatment with DMSO or CBX. (DMSO: 4 animals, 7 DRG, 26 cells CBX: 3 animals, 5 DRG, 34 cells) R) Depiction of targeted cellular processes for molecular screen. S) Quantification of the average number of Ca^2+^ events per DRG following pharmacological screen. (DMSO: 19 animals, 64 DRG, HMR1556: 5 animals 15 DRG Apyrase: 5 animals, 15 DRG Thaps: 4 animals, 13 DRG, GsMTx4: 5 animals, 13 DRG CBX: 5 animals, 14 DRG) Scale bar is 10μm (A,B,D,F,K,L,O). Ca^2+^ transient events are timepoints containing a z score of the integrated density of fluorescence greater than 2.58 (C,E,G,H,I,J,K,L,). Statistical tests: One-way ANOVA followed and represented by post hoc tukey test: (I,J,N,P), Unpaired student t tests: (Q), One-way Brown-Forsythe ANOVA followed and represented by post hoc dunnett test: (S), Correlation coefficient test: (M,N,Q).

**Figure 2:**
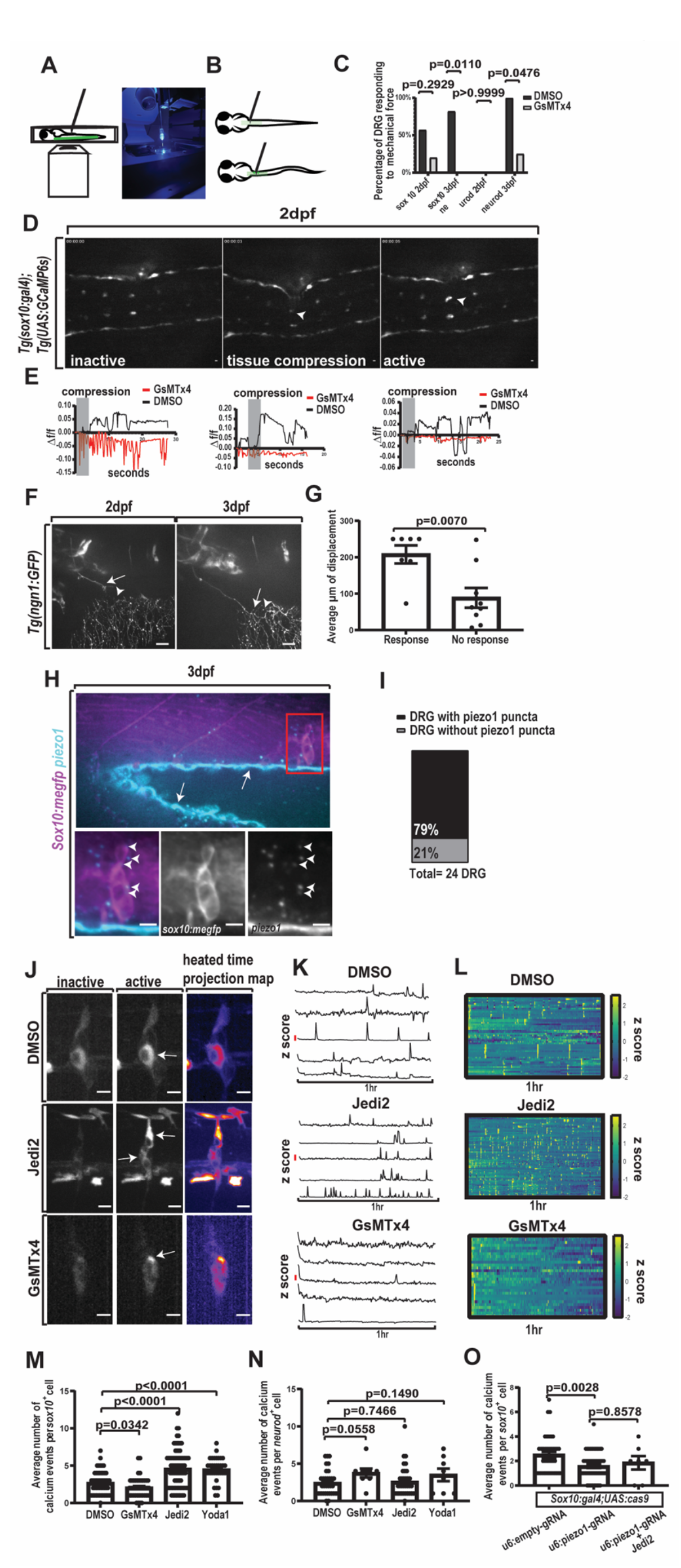
DRG are mechanosensitive and express *piezo1* A) LEFT Depiction of mechanical compression assay where animal is mounted dorsally on inverted spinning disk confocal with a dextran loaded microneedle mounted above the animal. RIGHT image of mechanical compression assay apparatus. B) Depiction of mechanical compression assay with needle placing force on DRG. C) Quantification of the percent of DRG responding to mechanical force in animals expressing *Tg(sox10:gal4+myl7); Tg(uas:GCaMP6s)* (labeled sox10) or expressing *Tg(neurod:gal4+myl7); Tg(uas:GcaMP6s)* (labeled neurod) and treated with either DMSO or GsMTx4 at both 2 and 3 dpf. (sox10 2dpf DMSO: 7 animals, 7 DRG, sox10 2dpf GsMTx4: 5 animals, 5 DRG neurod 2dpf DMSO: 5 animals 5 DRG neurod 2dpf GsMTx4: 4 animals, 4 DRG sox10 3dpf DMSO: 11 animals, 11 DRG sox10 3dpf GsMTx4: 4 animals, 4 DRG neurod 3dpf DMSO: 9 animals, 9 DRG neurod 3dpf GsMTx4: 4 animals, 4 DRG) D) Confocal image taken of the mechanical compression assay in 2 dpf animal expressing *Tg(sox10:gal4+myl7); Tg(uas:GcaMP6s)*. Images show an inactive time point, a time point with tissue compression, and an active time point in response to tissue compression. Inactive and active DRG marked with an arrow. E) Quantification of the change in integrated density of fluorescence of a DRG in a 2 dpf animal treated with either DMSO or GsMTx4. Change in integrated density of fluorescence is scored as time point subtracting the initial timepoint divided by time point(*Δf*/*f*). Grey box notes timepoints of tissue compression. Each graph is an example of this quantification from a biological replicate. F) Confocal z-projection of peripheral DRG axon in an animal expressing *Tg(ngn1:GFP)* at 2 and 3 dpf. Arrow notes the end processes of the peripheral axon. Arrowhead denotes peripheral axons from Rohon beard neurons. G) Average distance (μm) of DRG displacement needed to elicit a response. (16 animals, 16 DRG) H) Confocal images of RNAscope-*piezo1* and Immunohistochemistry-GFP in *Tg(sox10:meGFP)* animals. GFP is shown in magenta and *piezo1* is shown in cyan. Arrowheads indicate *piezo1* puncta. Arrows indicate autofluorescence. I) Quantification of DRG at 3 dpf with *piezo1* puncta and without *piezo1* puncta. (9 animals, 24 DRG) J) Confocal images of 3 dpf animals expressing *Tg(sox10:gal4+myl7); Tg(uas:GCaMP6s)*. Red colors indicate a higher intensity of fluorescence, and blue colors indicate a lower intensity of fluorescence. Images depicted are of animals either treated with 2% DMSO, 40μM Jedi2, or 1μM GsMTx4. Arrows note active cells. K) Line graphs of z score of integrated density of fluorescence for a 1hr time period in 3 dpf animals expressing *Tg(sox10:gal4+myl7); Tg(uas:GCaMP6s)* that were treated with either 2% DMSO, 40μM Jedi2, or 1μM GsMTx4. A z score greater than 2.58 indicates an active Ca^2+^ event. Red scale bar shows a z score of 2.58. L) Heatmaps of the z score of individual *sox10^+^* cells from animals in G-H during a 1 hr period of Ca^2+^ imaging. Yellow notes a high z score (2.58 or greater). (DMSO: 8 animals, 17 DRG, 52 cells, Jedi2: 7 animals, 18 DRG, 79 cells, GsMTx4: 6 animals, 16 DRG, 44 cells) M) Quantification of the average number of Ca^2+^ events per *sox10^+^* cell in animals treated with either 2% DMSO, 1μM GsMTx4, 100μM Yoda1, or 40μM Jedi2. (DMSO: 8 animals, 17 DRG, 52 cells, Jedi2: 7 animals, 18 DRG, 79 cells, GsMTx4: 6 animals, 16 DRG, 44 cells, Yoda1: 4 animals, 9 DRG, 35 cells) N) Quantification of the average number of Ca^2+^ events per *neurod^+^* cell in 3 dpf animals expressing *Tg(neurod:gal4+myl7); Tg(uas:GCaMP6s)* that were treated with either 2% DMSO, 1μM GsMTx4, 40μM Jedi2, or 100μM Yoda1. (DMSO: 10 animals, 24 DRG, 33 cells, Jedi2: 5 animals, 18 DRG, 35 cells, GsMTx4: 4 animals, 8 DRG, 8 cells, Yoda1: 4 animals, 7 DRG, 8 cells) O) Quantification of the average number of Ca^2+^ events per *sox10^+^* cell following genetic manipulation via injection of *uas:cas9mkate-u6:piezo1gRNA* or *uas:cas9mkate-u6:emptygRNA* into animals expressing *Tg(sox10:gal4+myl7); Tg(uas:GCaMP6s)* at 3dpf. Additionally a group of *Tg(sox10:gal4+myl7); Tg(uas:GCaMP6s)* injected with *uas:cas9mkate-u6:piezo1gRNA* and treated with 40μM Jedi2 treatment was also included in the experiment. (u6:emptygRNA: 6 animals, 16 DRG, 40 cells, u6:piezo1gRNA: 4 animals, 13 DRG, 57 cells, u6:piezo1gRNA+Jedi2: 4 animals, 4 DRG, 7 cells) Scale bar 10μm (D,F,H,J). Statistical tests: unpaired t test (G, M,N), Fisher’s exact (D).

Neural cells can exhibit distinct spontaneous Ca^2+^ transient events. To explore if DRG satellite glia exhibit distinct subtypes of Ca^2+^ transients, we created activity profiles for each cell in a given DRG from z-score calculations in 1 hour movies of *Tg(sox10:gal4+myl7); Tg(uas:GCaMP6s); Tg(neurod:tagRFP)* 3dpf animals. Using this data, we could then compare when each individual cell in a DRG was active compared to the other cells in the DRG. We found that individual *sox10^+^* cells displayed Ca^2+^ transients simultaneously with other *sox10^+^* cells in the DRG (Fig 1F), consistent with previous descriptions of simultaneous Ca^2+^ transients in glial networks.[11,39,40] However, we also identified a subset of Ca^2+^ transients that occurred in cells when no neighboring cell is active (Fig 1F). We define these Ca^2+^ transients in this report as isolated Ca^2+^ transients.

Calcium microdomains are also known to be present in several glial types[10,34,41]. Therefore, we tested if Ca^2+^ microdomains are also present in the DRG during development. To do this, we imaged animals expressing a membrane localized GCaMP6s by injecting *Tg(sox10:gal4+myl7)* embryos with *uas:GCaMP6s-caax* and imaging at 3 dpf. In order to initially capture and identify these quick dynamic events we imaged animals for a 10 minute period with 5 second intervals capturing the entire DRG. We defined Ca^2+^ microdomains as small regions with significant changes in integrated density of fluorescence of GCaMP6s-caax (Fig 1D,E). We quantified the duration of these microdomains and found that they lasted on average for 11.81+/-9.914 seconds (n=15 cells, 8 DRG, 7 animals) (Fig 1I). We also quantified the average volume of these microdomains and found that they were on average 30.10+/-18.51 μm^3^ (n=15 cells, 8 DRG, 7 animals) (Fig 1J). Together, these results indicate DRG satellite glia exhibit at least three distinct Ca^2+^ transient events during development: isolated, simultaneous, and microdomains.

### Satellite glia cell networks are established during early DRG construction

To understand how these types of activity may change over development, we quantified the average amount of isolated and simultaneous Ca^2+^ transient events in the same animal at 2, 3, and 4 dpf. While we did not see a significant change in isolated Ca^2+^ transients over this developmental period (2dpf: n=46 cells, 10 DRG, 6 animals, 3dpf: n=27 cells, 6 DRG, 4 animals, 4dpf: n=34 cells, 7 DRG, 4 animals) (Fig 1G), there was a noted significant increase in the number of simultaneous Ca^2+^ transient events after 2 dpf (2dpf vs. 3dpf: p=0.0019, 2dpf vs. 4dpf: p=0.0449 post hoc tukey test) (2dpf: n=46 cells, 10 DRG, 6 animals, 3dpf: n=27 cells, 6 DRG, 4 animals, 4dpf: n=34 cells, 7 DRG, 4 animals) (Fig 1H). This work indicates distinct developmental properties between isolated and simultaneous subtypes.

Current research proposes that DRG satellite glia form networks *in vitro*[42–44]. To further test if this occurs *in vivo* and to determine when in development it arises, we measured synchronized networks in *sox10^+^* cells. To identify a synchronized network of cells we compared the Ca^2+^ transient profiles of individual cells by computing the correlation between two Ca^2+^ transient profiles. To determine how this changed in development, we quantified the percent of high correlation coefficients (>0.5) per cell in each DRG at 2, 3, and 4 dpf. By creating network maps of individual DRG that show how the activity of each cell is related (Fig 1K-M), we found that *sox10^+^* cells at 2 dpf had an average of 37.48%±32.08% high correlation coefficients (n=50 cells, 10 DRG, 6 animals). At 3 dpf we measured that *sox10^+^* cells had an average of 39.33%±30.61% high correlation coefficients (n=27 cells, 6 DRG, 4 animals) and by 4 dpf, *sox10^+^* cells had a significant increase in the percent of high correlation coefficients, with an average of 58.26%±32.13% high correlation coefficients (n=34 cells, 7 DRG, 4 animals) (2dpf vs. 4dpf: p=0.0044 post hoc tukey test, 2dpf n=46 cells, 3dpf n=27 cells, 4dpf n=34 cells) (Fig 1N). Additionally, we observed that the percent of cells displaying Ca^2+^ transients together increased by 4 dpf (Sup Fig3A-C). These data are consistent with the hypothesis that DRG satellite glial networks are present *in vivo* and form by at least the third day of DRG construction in zebrafish.

If glial networks are forming, we hypothesized that gap junctions may also increase during the time when synchronized Ca^2+^ transients are present. Cxn43 is known to be present in satellite glia and contribute to gap junctions in synchronized neural networks[45–47]. Therefore, we stained for Cxn43 at 2, 3, and 4 dpf in animals expressing *Tg(sox10:meGFP),* which labels satellite glia in the DRG with membrane-localized GFP. The 2 dpf DRG had an average of 0.500±0.707 Cxn43 puncta (n=18 DRG, 6 animals). This increased to an average of 1.000±0.845 Cxn43 puncta per DRG at 3 dpf (n=29 DRG, 10 animals) and by 4 dpf, there was a significant increase in the number of Cxn43 puncta present in the DRG with an average of 2.208±1.062 Cxn43 puncta per DRG (n=24 DRG, 8 animals) (2dpf vs. 4dpf: p<0.0001, 3dpf vs 4dpf: p<0.0001 post hoc tukey test) (Fig 1O, 1P). These results support the hypothesis that DRG cells begin forming glial connections during its earliest construction.

To determine if there are functional gap junction connections, we treated animals expressing *Tg(sox10:gal4+myl7); Tg(uas:GCaMP6s); Tg(neurod:tagRFP)* with either Carbenoxolone (CBX), a gap junction inhibitor, or a control treatment of DMSO. Animals treated with CBX at 3 dpf demonstrated a significant decrease in the percent of high correlation coefficients with an average percent of 22.00%±22.34% (n=34 cells, 5 DRG, 3 animals) compared to an average percent of high correlation coefficients of 36.58%±31.22% when treated with a DMSO control (n=26 cells, 7 DRG, 4 animals) (DMSO vs. CBX: p=0.0391 unpaired t test) (Fig 1Q). These results strongly support the idea that functional gap junctions are present in glial networks in DRG during its early construction.

### Satellite glia Ca^2+^ transients are impacted by altering mechanobiology

Our measurements indicated that DRG satellite glia cells demonstrate distinct Ca^2+^ transients. To identify potential molecular components involved in these Ca^2+^ transients, we performed a chemical screen targeting various chemical signals shown to affect Ca^2+^ transients using transgenic animals expressing *Tg(sox10:gal4+myl7); Tg(uas:GCaMP6s); Tg(neurod:tagRFP)* (Fig 1R-S). Additionally, we included a broad-mechanosensitive ion channel antagonist, GsMTx4, because of the underappreciated role that mechanobiology has during neurodevelopment. We hypothesized that GsMTx4 would reduce the amount of observed Ca^2+^ transients if mechanobiology had an important role during early development. Each animal was exposed to the pharmacological agent 30 minutes prior and during the imaging window and then GCaMP6s intensity was measured for 1 hour with a 15s imaging interval. We reasoned that an overall change in the abundance of Ca^2+^ transients could help us identify molecules that are important for either isolated or simultaneous spontaneous Ca^2+^ transients. We found that GsMTx4 significantly reduced the amount of Ca^2+^ transients observed compared to DMSO (Fig 1S) (DMSO vs. GsMTx4: p<0.0001 post hoc Dunnett test). We also measured a significant change following treatment with Thapsigargin (Thaps) (Fig 1S) (DMSO vs. Thaps: p=0.0010 post hoc Dunnett test). While chemical signaling has been widely described in spontaneous Ca^2+^ transients, the role of mechanobiology in the process is less known, which led us to investigate the potential role of mechanobiology in spontaneous Ca^2+^ transients in the DRG.

To first explore the possibility that mechanical features impact spontaneous Ca^2+^ transients in the DRG, we tested if the cells in the developing DRG are sensitive to mechanical perturbation. To do this, we imaged the DRG of transgenic zebrafish expressing GCaMP6s in satellite glial *Tg(sox10:gal4+myl7); Tg(uas:GCaMP6s)* cells during tissue compression (Fig 2A). Tissue compression was administered by bending the animal with a microneedle as they were imaged on the confocal microscope (Fig 2B). At 2 dpf, 57% of DRG expressing *Tg(sox10:gal4+myl7); Tg(uas:GCaMP6s)* responded to tissue compression (n=7 DRG, 7 animals) (Fig 2C-D). We found that the average distance that a DRG moved with animal bending in order to elicit a response via GCaMP6s was 207.8 μm (n=16 DRG, 16 fish) (Fig 2G). It is possible that this response of *sox10^+^* cells was secondary to neuronal firing. We, therefore, tested if neurons fired in response to compression at 2 dpf in *Tg(neurod:gal4+myl7); Tg(uas:GCaMP6s)* animals but could not detect Ca^2+^ transients in neurons after compression (n=5 DRG, 5 animals) (Fig 2C). Examining DRG axonal projections in *Tg(ngn1:GFP)* animals also showed that neurons at 2 dpf did not have peripheral axons at their final targets in the periphery. It therefore seems unlikely that such Ca^2+^ transients in *sox10^+^* satellite glia after tissue compression are secondary to neuronal activity. To understand if *sox10^+^* satellite glia continued to be sensitive to mechanical compression, we repeated this assay at 3 dpf. By 3 dpf, 82% (n=11 DRG, 11 animals) of DRG expressing *Tg(sox10:gal4+myl7); Tg(uas:GCaMP6s)* responded to tissue compression (Fig2C). At 3 dpf, 100% (n=5 DRG, 5 animals) of DRG expressing *Tg(neurod:gal4+myl7); Tg(uas:GCaMP6s)* also demonstrated Ca^2+^ transients after tissue compression (Fig 2C). While the neuronal population of the DRG does respond to tissue compression at a later age, our data suggests satellite glia respond to the mechanical tissue compression at early ages without neuronal activation.

If this response to mechanical force is mediated by mechanosensitive ion channels, we would hypothesize that it would be reduced upon treatment of GsMTx4, which broadly blocks mechanosensitive ion channels. To test this hypothesis, we imaged animals expressing either *Tg(sox10:gal4+myl7); Tg(uas:GCaMP6s); Tg(neurod:tagRFP)* or *Tg(neurod:gal4+myl7); Tg(uas:GCaMP6s)* that were treated with GsMTx4. We found that treatment with GsMTx4 reduced the response to mechanical stimuli to 20% of animals (n=5 DRG, 5 animals) expressing *sox10^+^* GCaMP6s at 2 dpf (Fig 2C,E). At 3 dpf when treated with GsMTx4 there was a significant reduction in response with 0% of animals (n=4 DRG, 4 animals) expressing *sox10*^+^ GCaMP6s responded to mechanical force and 25% of animals (n=4 DRG, 4 animals) expressing neuronal GCaMP6s responded to mechanical force (*sox10* 3dpf DMSO vs. GsMTx4: p=0.0110, neurod 3dpf DMSO vs. GsMTx4: p=0.0476 Fisher’s exact test) (Fig 2C). These data support the hypothesis that DRG are responsive to mechanical forces and identify that *sox10^+^* cells are mechanosensitive, at least partially independent of neuronal activity.

### Satellite glia Ca^2+^ transients can be altered by manipulating Piezo1

We next explored the potential molecular determinant of this mechanical component. The mature DRG is known to express mechanosensitive channels Piezo1 and Piezo2, however Piezo2 is restricted to neurons while Piezo1 is expressed in neurons and satellite glia in mice[31]. To investigate this in zebrafish, we utilized RNAscope to determine spatiotemporal distribution of *piezo1* RNA in animals expressing *Tg(sox10:meGFP)*. We found that 79% of DRG (n=24 DRG, 9 animals) at 3 dpf contained *piezo1* RNAscope puncta within *sox10^+^* satellite glia (Fig 2H,I). Additionally we utilized Whole-mount HCR-FISH targeting *piezo1* RNA 3dpf animals expressing *Tg(sox10:meGFP)* and found similar expression of *piezo1* (Sup Fig 4).

**Figure 4:**
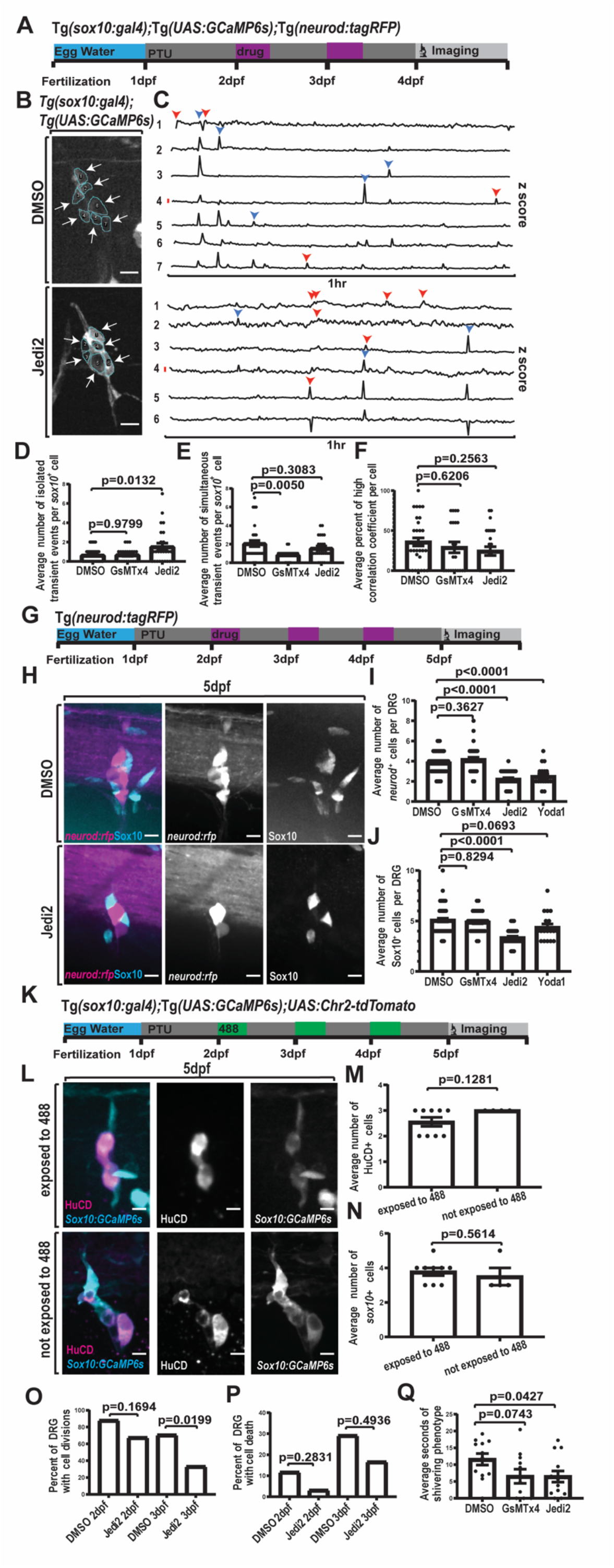
Increased isolated activity via Piezo1 decreases DRG cell divisions and reduces response to cold stimulus A) Timeline of experimental process where animals were treated with either 2% DMSO, 1μM GsMTx4, or 40μM Jedi2 for 30 minutes each day at 2 and 3 dpf. B) Confocal z-projection of DRG in animals expressing *Tg(sox10:gal4+myl7); Tg(uas:GCaMP6s)* following consecutive days of treatment with either DMSO or Jedi2. ROIs are traced individual cells labeled with a number. Arrows denote active cells. C) Line graphs of the z score of the integrated density of fluorescence over a 1 hr period. Numbers correlate with ROIs in (B). Red line signifies a z score of 2.58. Blue arrowheads note simultaneous active time points. Red arrowheads note isolated active time points. D-F) Quantification of the average number of isolated (D), simultaneous Ca^2+^ activity events per *sox10^+^* Ils (E) or average percent of high correlation coefficient(F) per *sox10^+^* in 4 dpf animals expressing *Tg(sox10:gal4+myl7); Tg(uas:GCaMP6s)* that were treated at 2 and 3 dpf with DMSO, GsMTx4, or Jedi2. (DMSO: 4 animals, 6 DRG, 32 cells, GsMTx4: 3 animals, 4 DRG, 22 cells, Jedi2: 4 animals, 5 DRG, 25 cells) G) Timeline of experimental process where animals were treated with either 2% DMSO, 1μM GsMTx4, 100μM Yoda1, or 40μM Jedi2 for 30 minutes each day at 2-4 dpf. Animals were then fixed and processed for imaging at 5 dpf. H) Confocal z-projections of 5 dpf DRG in animals expressing *Tg(neurod:tagRFP)* and stained for Sox10. Magenta displays *Tg(neurod:tagRFP)*, and cyan displays Sox10. I-J) Quantification of the average number of *neurod^+^* (I) and Sox10 (J) cells in 5 dpf animals treated with either 2% DMSO, 1μM GsMTx4, 40μM Jedi2, or 100μM Yoda1 for 30 minutes each day from 2-4 dpf. (DMSO: 15 animals, 53 DRG, GsMTx4: 10 animals, 34 DRG, Jedi2: 10 animals, 39 DRG, Yoda1: 7 animals, 18 DRG) K) Timeline of experimental process where animals expressing *Tg(sox10:gal4+myl7); Tg(uas:GCamp6s); uas:Chr2-tdTomato* were exposed to 488nm light for 30 minutes each day during targeted developmental period. Animals were fixed, stained, and processed for imaging at 5dpf. L) Confocal z-projections of 5dpf DRG in animals expressing *Tg(sox10:gal4+myl7); Tg(uas:GCamp6s); uas:Chr2-tdTomato* and stained for HuCD expression. Magenta displays HuCD^+^ neurons. Cyan displays *sox10^+^* satellite glia. (M-N) Quantification of the average number of HuCD^+^ neurons (M) and *sox10^+^* satellite glia (N) in 5dpf animals expressing *Tg(sox10:gal4+myl7); Tg(uas:GCaMP6s); uas:Chr2-tdTomato* and either exposed or not exposed to 488nm light for 30 minutes each day during development. (exposed: 4 animals, 9 DRG, not exposed: 4 animals, 4 DRG) O-Q) Quantification of the percent of DRG with cell divisions (O) and/or cell deaths (P) in 24-hour time lapses of *Tg(sox10:meGFP); Tg(neurod:tagRFP)* animals treated either with 2% DMSO or 40μM Jedi2 from 2-3 dpf or 2-4 dpf. (2dpf DMSO: 6 animals, 17 DRG, 2dpf Jedi2: 8 animals, 31 DRG, 3dpf DMSO: 8 animals, 24 DRG, 3dpf Jedi2: 8 animals, 24 DRG) Q) Quantification of the average duration (seconds) of shivering in of 5 dpf animals that were treated with 2% DMSO, 1μM GsMTx4, or 40μM Jedi2 for 30 min each day from 2-4 dpf. (DMSO: 12 animals, GsMTx4: 11 animals, Jedi2: 13 animals) Scale bar is 10μm (B,H,L). Statistical tests: one-way ANOVA followed and represented by post hoc dunnett test (D, E, F, I, J), Fisher’s exact: (O, P), unpaired t test: (M,N,Q).

To test if DRG contain functional Piezo1, we treated animals expressing *Tg(sox10:gal4+myl7); Tg(uas:GCaMP6s); Tg(neurod:tagRFP)* with Yoda1 and Jedi2, known Piezo1 specific agonists (Fig 2J-L)[48,49]. We found the average amount of Ca^2+^ transients per *sox10^+^* cell in a 1 hour timelapse in DMSO controls was 2.69±1.55 Ca^2+^ transient events, which was significantly less than the average 4.48±1.58 Ca^2+^ transient events or 4.58±2.54 Ca^2+^ transient events observed when treated with Yoda1 or Jedi2 respectively (DMSO n=52 cells, 17 DRG, 8 animals, Yoda1 n=33 cells, 9 DRG, 4 animals, Jedi2 n=79 cells, 18 DRG, 7 animals) (DMSO vs. Yoda1: p<0.0001 unpaired t test, DMSO vs. Jedi2: p<0.0001 unpaired t test) (Fig 2M). Furthermore, we repeated this assay treating with GsMTx4 to investigate whether inhibiting mechanosensitive ion channels reduces spontaneous Ca^2+^ transients. This treatment significantly reduced the average amount of Ca^2+^ transients per *sox10^+^* cell to an average of 2.05±1.36 Ca^2+^ transient events (n=44 cells, 16 DRG, 6 animals) (DMSO vs GsMTx4: p=0.0342 unpaired t test) (Fig 2 J-M).

One possible explanation for an increase in Ca^2+^ transients in *sox10^+^* satellite glia is that the *sox10^+^* satellite glia are active in response to neuronal activity. To investigate whether the observed change in Ca^2+^ transients in *sox10^+^* satellite glia was a consequence of altered neuronal activity, we treated animals expressing *Tg(neurod:gal4+myl7); Tg(uas:GCaMP6s)* with Piezo1 agonists and quantified the amount of Ca^2+^ transients per DRG neuron. We found following DMSO treatment that DRG neurons exhibited an average of 2.42±1.71 Ca^2+^ transient events per hour. When animals were treated with either Yoda1 or Jedi2, an average of 3.50±2.39 or 2.57±2.00 Ca^2+^ transient events per hour respectively could be detected. When animals were treated with GsMTx4, there was an observed 3.75±1.67 average number of Ca^2+^ transient events (DMSO n=33 neurons, 24 DRG, 10 animals, Yoda1 n=8 neurons, 7 DRG, 4 animals, Jedi2 n=35 neurons, 18 DRG, 5 animals, GsMTx4 n=8 neurons, 8 DRG, 4 animals) (Fig 2N). Overall, we found that Piezo1 agonists did not contribute to an increase in Ca^2+^ transients in the *neurod^+^* population. These data are most consistent with the hypothesis that *sox10^+^* satellite glia display Ca^2+^ transients in response to Piezo1 agonists independent of an increase in neuronal activity.

In addition to these pharmacological treatments we also sought to do a genetic manipulation to identify if the endogenous Piezo1-mediated Ca^2+^ transient was present in satellite glia. We utilized a *uas:cas9mkate-u6:piezo1gRNA* construct designed with a *piezo1* gRNA to knockout *piezo1* in satellite glia.[50] This construct was injected into animals expressing *Tg(sox10:gal4); Tg(uas:GCaMP6s)*. We then imaged animals expressing *Tg(sox10:gal4); Tg(uas:GCaMP6s)*; *uas:cas9mkate-u6:piezo1gRNA* and quantified the average amount of Ca^2+^ transients in comparison with animals injected with *uas:cas9mkate-u6:emptygRNA*, a construct containing an empty gRNA cassette. We found in animals injected with *uas:cas9mkate-u6:piezo1gRNA* there was an average of 1.579+/− 1.117 Ca^2+^ transients (n=57 cells, 13 DRG, 4 animals). This was significantly lower than the average amount of Ca^2+^ transients observed in *uas:cas9mkate-u6:emptygRNA* injected animals where there was an average of 2.500+/-1.536 Ca^2+^ transient events (n=40 cells, 16 DRG, 6 animals) (p = 0.0028, post hoc tukey test) (Fig 2O). Additionally we injected animals expressing *Tg(sox10:gal4); Tg(uas:GCaMP6s)* with *uas:cas9mkate-u6:piezo1gRNA* and treated the animals Jedi2. These animals had on average 1.857+/-1.464 Ca^2+^ transients (n=7 cells, 4 DRG, 4 animals) which was not significantly different from untreated *uas:cas9mkate-u6:piezo1gRNA* injected animals (p = 0.8578, post hoc tukey test) (Fig 2O), supporting that this manipulation is specific to Piezo1. Together, these findings support the idea that *piezo1* contributes to spontaneous Ca^2+^ transients that are observed.

We demonstrated that DRG satellite glial cells have distinct Ca^2+^ transients (Fig 1F-H) but the underlying mechanism of those transients is unknown. We therefore tested if subtypes of Ca^2+^ transients were differentially modulated by Piezo1. To investigate the role of Piezo1 in isolated and simultaneous Ca^2+^ transient events, we quantified the effect of Yoda1, Jedi2, GsMTx4, and DMSO treatment on these Ca^2+^ transient events. In DMSO-treated animals, *sox10^+^* cells displayed an average 0.46±0.78 isolated Ca^2+^ transient events. When treated with Yoda1 or Jedi2, *sox10^+^* cells displayed an average of 2.28±1.72 or an average of 1.31±1.22 isolated Ca^2+^ transient events respectively. We found that when treated with GsMTx4, *sox10^+^* cells displayed an average of 0.91±1.2 isolated Ca^2+^ transient events (DMSO n=24 cells, 5 DRG, 4 animals, Yoda1 n=28 cells, 6 DRG, 3 animals, Jedi2 n=59 cells, 18 DRG, 7 animals, GsMTx4 n=23 cells, 5 DRG, 3 animals) (Fig 3A-C). These results show that Piezo1 agonists significantly increased the amount of isolated Ca^2+^ transients (DMSO vs. Yoda1: p<0.0001, DMSO vs. Jedi2: p=0.0024 unpaired t tests), while mechanosensitive antagonists did not significantly decrease isolated Ca^2+^ transients (DMSO vs GsMTx4: p=0.1294 unpaired t test) (Fig 3C). In contrast, simultaneous Ca^2+^ transient events were not significantly different in Yoda1 or Jedi2-treated animals compared to controls (DMSO n=24 cells, 5 DRG, 4 animals, Yoda1 n=28 cells, 6 DRG, 3 animals, Jedi2 n=59 cells, 18 DRG, 7 animals, GsMTx4 n=23 cells, 5 DRG, 3 animals) (Fig 3D). In the case of mechanosensitive antagonists, animals displayed a significant decrease in the number of simultaneous Ca^2+^ transient events compared to DMSO control when treated with GsMTx4 (DMSO vs. GsMTx4: p=0.0003 unpaired t test).

**Figure 3:**
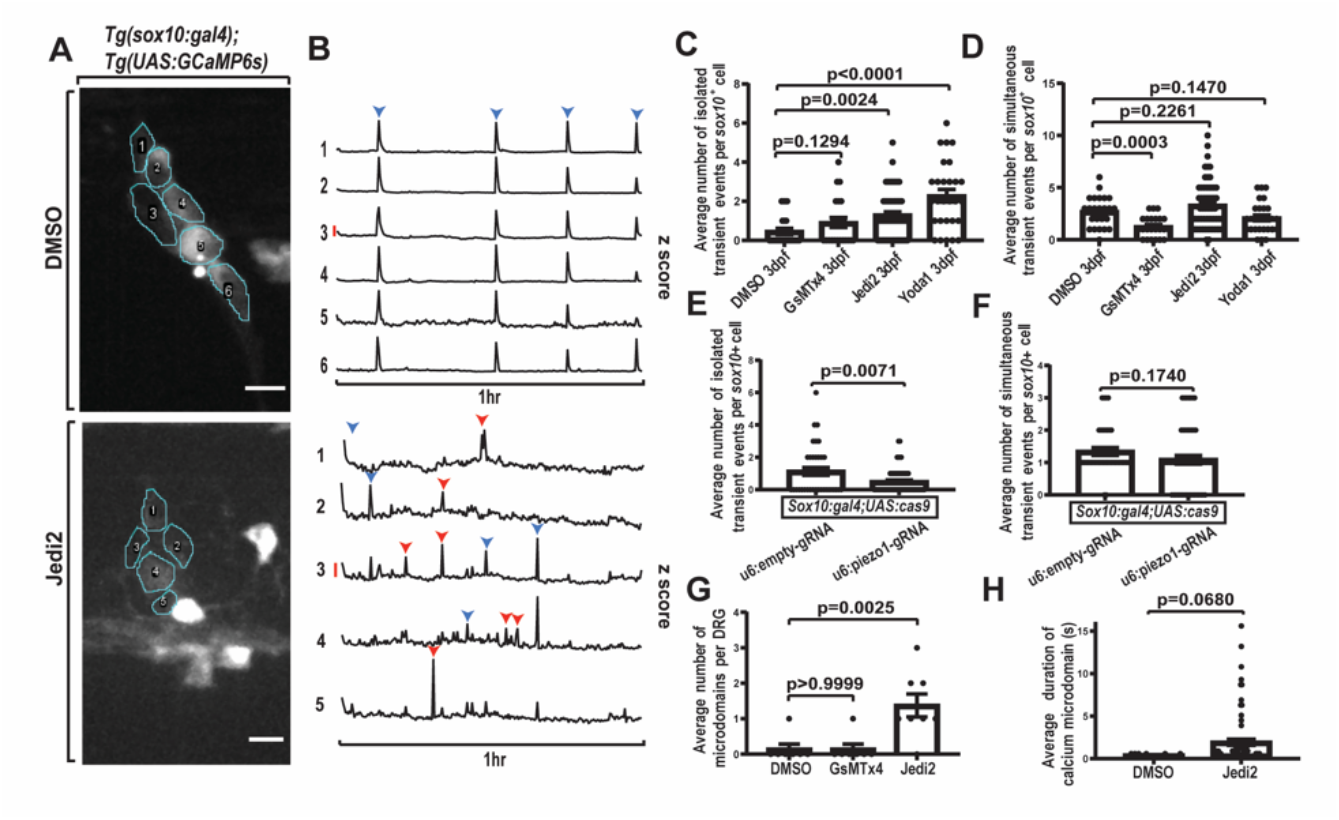
Piezo1 overactivation increases isolated calcium activity in DRG A) Confocal z-projection of DRG in 3 dpf animal expressing *Tg(sox10:gal4+myl7); Tg(uas:GCaMP6s)* **and treated with either 2% DMSO or 40μM Jedi2. Individual cells are traced for ROIs and labeled with a number. B) Line graphs of the z score of the integrated density of fluorescence over a 1 hr period. Each numbered line graph corresponds to a numbered ROI. Red scale bar represents a z score of 2.58. Blue arrowheads note simultaneously active time points. Red arrowheads note isolated active time points. C) Quantification of the average number of isolated Ca^2+^ transient events per** *sox10^+^* **cell in 3 dpf animals expressing** *Tg(sox10:gal4+myl7); Tg(uas:GCaMP6s)* **that were treated with either 2% DMSO, 1μM GsMTx4, 100μM Yoda1, or 40μM Jedi2. (DMSO: 4 animals, 5 DRG, 24 cells, GsMTx4: 3 animals, 5 DRG, 23 cells, Jedi2: 7 animals, 18 DRG, 59 cells, Yoda1: 3 animals, 6 DRG, 28 cells) D) Quantification of the average number of simultaneous Ca^2+^ transient events per** *sox10^+^* **cell in 3 dpf animals expressing** *Tg(sox10:gal4+myl7); Tg(uas:GCaMP6s)* **that were treated with either 2% DMSO, 1μM GsMTx4, 100μM Yoda1, or 40μM Jedi2. (DMSO: 4 animals, 5 DRG, 24 cells, GsMTx4: 3 animals, 5 DRG, 23 cells, Jedi2: 7 animals, 18 DRG, 59 cells, Yoda1: 3 animals, 6 DRG, 28 cells) E) Quantification of the average number of isolated Ca^2+^ transient events following genetic manipulation via CRISPR/Cas9 targeting** *piezo1* **or empty gRNA cassette in 3dpf animals expressing** *Tg(sox10:gal4+myl7); Tg(uas:GCaMP6s) uas:cas9mkate-u6:piezo1gRNA* **or** *uas:cas9mkate-u6:emptygRNA***. (u6:emptygRNA: 6 animals, 14 DRG, 38 cells, u6:piezo1gRNA: 4 animals, 13 DRG, 57 cells) F) Quantification of the average number of simultaneous Ca^2+^ transient events following genetic manipulation via CRISPR/Cas9 targeting** *piezo1* **or empty gRNA cassette in 3dpf animals expressing** *Tg(sox10:gal4+myl7); Tg(uas:GCaMP6s) uas:cas9mkate-u6:piezo1gRNA* **or** *uas:cas9mkate-u6:emptrygRNA***. (u6:emptygRNA: 6 animals, 14 DRG, 38 cells, u6:piezo1gRNA: 4 animals, 13 DRG, 57 cells) G) Quantification of the average number of microdomains per DRG in 3 dpf animals expressing** *Tg(sox10:gal4+myl7); uas:GCaMP6s-caax* **that were treated with either 2% DMSO, 1μM GsMTx4, or 40μM Jedi2 for 30 minutes prior to imaging. (DMSO: 7 animals, 7 DRG, GsMTx4: 7 animals, 7 DRG, Jedi2: 6 animals, 8 DRG) H) Quantification of the average duration of microdomains in 3 dpf animals expressing** *Tg(sox10:gal4+myl7); uas:GCaMP6s-caax* **that were treated with either 2% DMSO or 40μM Jedi2 for 30 minutes prior to imaging. (DMSO: 8 animals, 8 DRG, Jedi2: 5 animals, 7 DRG) Scale bar is 10μm (A). Statistical tests: unpaired t test (C, D,E,F,H), one-way ANOVA followed and represented by post hoc dunnett test(G).**

To complement the pharmacological approach, we also quantified these distinct Ca^2+^ transient events in genetic manipulations following the injection of *uas:cas9mkate-u6:piezo1gRNA* and *uas:cas9mkate-u6:emptygRNA*. Animals injected with the *empty-gRNA* had an average of 1.132+/-1.474 isolated Ca^2+^ transient events and an average of 1.342+/-0.7453 simultaneous Ca^2+^ transient events (n=38 cells, 14 DRG, 6 animals) (Fig 3E-F). Animals injected with the *piezo1-gRNA* had an average of 0.4912+/-0.7820 isolated Ca^2+^ transient events and an average of 1.088+/− 0.9688 simultaneous Ca^2+^ transient events (n=57 cells, 13 DRG, 4 animals) (Fig 3E-F). We found there was a significant reduction in the number of isolated Ca^2+^ transient events in animals injected with *uas:cas9mkate-u6:piezo1gRNA* compared to *uas:cas9mkate-u6:emptygRNA* (p = 0.0071, unpaired t test). The average number of simultaneous Ca^2+^ transient events were not significantly different following these manipulations (p=0.1740, unpaired t test). Together, the pharmacological and genetic manipulations support the hypothesis that Piezo1 contributes specifically to isolated Ca^2+^ transients.

We also quantified the number of Ca^2+^ microdomains following manipulations of Piezo1 in *Tg(sox10:gal4+myl7)* animals injected with *uas:GCaMP6s-caax*. In DMSO-treated animals, *sox10^+^* cells displayed an average number of 0.14±0.38 Ca^2+^ microdomains. Animals treated with Jedi2 displayed an average number of 1.38±0.92 Ca^2+^ microdomains. When animals were treated with GsMTx4 we observed an average of 0.14±0.38 Ca^2+^ microdomains (DMSO n=7 DRG, 7 animals, Jedi2 n=8 DRG, 6 animals, GsMTx4 n=7 DRG, 7 animals) (Fig 3G). Similar to isolated Ca^2+^ transients, we found a significant increase in the average number of Ca^2+^ microdomains per DRG when animals were treated with a Piezo1 agonist (DMSO vs. Jedi2: p=0.0025 post hoc dunnett test) (Fig 3G). We also quantified the average duration of the identified Ca^2+^ microdomains and did not find a significant difference (Fig 3H). These additional findings suggest that Piezo1-mediated mechanical forces contribute to the number of observable Ca^2+^ microdomain events in addition to isolated Ca^2+^ transient events.

### Altering Piezo1 has functional consequences to DRG development

The specific function of isolated Ca^2+^ transients is relatively unknown. We therefore used Piezo1 manipulations to test the potential functional consequence of increasing isolated Ca^2+^ transients. We first hypothesized that Piezo1-sensitive isolated Ca^2+^ transients were important for the formation of synchronized glial networks. These glial networks form between 2-4 dpf (Figure 1N). Therefore, to first test this hypothesis, we treated animals expressing *Tg(sox10:gal4+myl7); Tg(uas:GCaMP6s); Tg(neurod:tagRFP)* with either DMSO, GsMTx4, or Jedi2 for 30 minutes daily at 2 and 3 dpf. We performed our Ca^2+^ imaging paradigm on these animals at 4 dpf and first assessed the amount of isolated and spontaneous Ca^2+^ transient events following consecutive days of treatment. Following consecutive days treated with DMSO, *sox10^+^* cells showed an average of 2.06±1.85 simultaneous Ca^2+^ transients and an average of 0.63±0.75 isolated Ca^2+^ transients. Following treatment on consecutive days with Jedi2, *sox10^+^* cells showed an average of 1.56±1.19 simultaneous Ca^2+^ transients and an average of 1.52±1.85 isolated Ca^2+^ transients. Taken together these data confirm that following consecutive days of treatment, Jedi2 significantly increases the amount of isolated Ca^2+^ transients (DMSO vs. Jedi2: p=0.0132 post hoc dunnett test). Interestingly, after treatment with the broad-mechanosensitive antagonist, GsMTx4, on consecutive days in development, *sox10^+^* cells displayed an average of 0.86±0.56 simultaneous Ca^2+^ transient events and an average of 0.68±0.72 isolated Ca^2+^ transient events, significantly reducing the amount of simultaneous Ca^2+^ transients (DMSO vs. GsMTx4: p=0.0050 post hoc dunnett test) (DMSO n=32 cells, 6 DRG, 4 animals, Jedi2 n=25 cells, 5 DRG, 4 animals, GsMTx4 n=22 cells, 4 DRG, 3 animals) (Fig 4D,E). To answer whether these changes ultimately impacted synchrony, we quantified the average percent of high correlation coefficients per *sox10^+^* cell following these treatment paradigms. We found that when treated with DMSO, *sox10^+^* cells had an average of 35.88±28.78% of high correlation coefficients. When treated with either GsMTx4 or Jedi2, *sox10^+^* cells had an average of 29.09±31.65% high correlation coefficients or an average of 24.48±27.16% high correlation coefficients respectively (DMSO n=32 cells, 6 DRG, 4 animals, GsMTx4 n=22 cells, 4 DRG, 3 animals, Jedi2 n=25 cells, 5 DRG, 5 animals) (Fig 4F). There was no significant change in the average percent of high correlation coefficients when treated with Jedi2 or GsMTx4 and thereby inconsistent with the idea that Piezo1-sensitive isolated Ca^2+^ transients or Ca^2+^ microdomains impact the formation of a presumptive glial network, at least as identified by synchronous activity.

In addition to the synchronous glial networks that we identify form by 4 dpf, it is also known that the DRG rapidly expands during this developmental time[38]. We, therefore, tested the hypothesis that Piezo1-sensitive events like isolated Ca^2+^ transients could be important for DRG expansion. To do this, we increased isolated Ca^2+^ transients via Piezo1 through Jedi2 or Yoda1 treatment and assessed the number of cells present in the DRG at 5 dpf. To assay both neuronal and glial expansion, we treated zebrafish expressing *Tg(neurod:tagRFP)* with Piezo1 agonists or a mechanosensitive antagonist for 30 minutes each day from 2-4 dpf, and then used immunohistochemistry against Sox10, thereby identifying TagRFP^+^ neurons and Sox10^+^ satellite glia at 5 dpf. Animals treated with DMSO on consecutive days displayed an average number of 3.85±1.05 *neurod^+^* cells and an average number of 5.09±1.33 Sox10^+^ cells. When animals were treated with GsMTx4, DRG also contained an average of 4.18±1.22 *neurod^+^* cells and 4.91±0.99 Sox10^+^ cells. However, treatment with Jedi2, resulted in DRG with an average of 2.23±0.81 *neurod^+^* cells and average number of 3.36±0.81 Sox10^+^ cells (Fig 4I-J), showing a reduction in the amount of DRG cells (DMSO vs. Jedi2: p<0.0001 post hoc dunnett test) (DMSO n=53 DRG, 15 animals, GsMTx4 n=34 DRG, 10 animals, Jedi2 n=39 DRG, 10 animals). With treatment with Yoda1, DRG (n=18 DRG, 7 animals) also displayed a significant reduction in DRG cells, with an average number of 2.50±1.04 *neurod^+^* cells (DMSO vs. Yoda1: p<0.0001, post hoc dunnett test). Taken together, these findings identify that altering Piezo1-sensitive isolated or microdomain Ca^2+^ transients impacts DRG development through a reduction in the number of cells present.

To investigate whether this change in cell abundance was a consequence of increased Ca^2+^ transients non-specific to Piezo1-mediated activity, we utilized an optogenetic approach to increase Ca^2+^ transients in the DRG. We injected *uas:Chr2-tdTomato* into animals expressing *Tg(sox10:gal4); Tg(uas:GCaMP6s)* and exposed the animals to 488nm light for 30 minutes each day from 2dpf until 4dpf (Fig 4K).[51] Prior to this experiment, we verified that the *uas:Chr2-tdTomato* injected construct would increase Ca^2+^ transients by exposing animals over a short period of time to 488nm light and quantifying the change in fluorescent intensity (Sup Fig 5). Following the 488nm exposure from 2-4dpf, we fixed the animals at 5dpf and stained for the neuronal marker HuCD (Fig 4L). We then quantified the number of HuCD^+^ cells and *sox10^+^* only cells to compare the number of both populations to a group of injected animals that were not exposed to 488nm light during development. In animals exposed to 488nm light we found an average of 2.556+/-0.527 HuCD^+^ neurons and an average of 3.778+/-0.667 *sox10^+^* satellite glia (n=9 DRG, 4 animals). The animals that were not exposed to 488nm light during development had an average number of 3.000 HuCD^+^ neurons and an average of 3.500+/-1.000 *sox10^+^* satellite glia (n=4 DRG, 4 animals). We found no significant difference in the number of neurons or satellite glia present in the DRG following after *Chr2* manipulation (Fig 4K-N). These findings support the hypothesis that increasing Piezo1-mediated isolated Ca^2+^ transients impacts the DRG via lowering cell abundance.

It is possible that this decrease in cell abundance was caused by a decrease in cell divisions or from an increase in cell death. In order to understand the cause of the decrease in cell abundance, we assessed the number of cell divisions and cell death occurring following consecutive treatment with Piezo1 agonists. To do this, we treated animals expressing *Tg(sox10:meGFP)* from 2-4 dpf with Jedi2 or a DMSO control. We then utilized overnight time lapse imaging to assay cell divisions or cell death. When treated with DMSO at 2 dpf, 88.24% of DRG had cell divisions (n=17 DRG, 6 animals). At 3 dpf following DMSO treatment, cell divisions occurred in 70.83% of DRG (n=24 DRG, 8 animals). Following treatment of Jedi2 at 2 dpf, 67.74% of DRG had cell divisions (n=31 DRG, 8 animals). At 3 dpf following consecutive treatment with Jedi2, 33.33% of DRG had cell divisions (n=24 DRG, 8 animals) (Fig 4O). We found a significant decrease in the number of cell divisions following consecutive treatment of Jedi2 at 2 and 3 dpf (DMSO vs Jedi2 3dpf: p=0.0199 Fisher’s exact test) (Fig 4O). However, we did not observe a significant change in the number of observed cell deaths (Fig 4P). The most likely explanation for this data is that the decrease in cell abundance in Piezo1-manipulated animals is from a reduction in cell divisions.

Lastly, we questioned whether this Piezo1-mediated decrease in cell abundance had functional consequences to the animal’s physiology. To answer this question, we treated animals with Piezo1 agonists and mechanical-channel antagonists during development and then tested the animals response to sensory stimuli. We previously demonstrated that larval zebrafish DRG neurons are active after zebrafish larvae are submerged in 4°C water[52,53]. This submersion causes a shivering phenotype that is at least partially dependent on intact DRG neurons and axons[52,53]. We treated animals with Jedi2 or GsMTx4 for 30 minutes each day from 2-4 dpf and then assayed sensory responses at 5 dpf. Following consecutive days of treatment, DMSO control treated animals had a shivering phenotype average duration of 11.60±5.9s. When treated on consecutive days with GsMTx4, animals had an average length of shivering of 6.55±6.97s. Following consecutive days of treatment with Jedi2, animals had an average length of shivering of 6.47±6.01s. There was a significant decrease in the length of shivering following consecutive days of Jedi2 treatment suggesting that overactivation of Piezo1 during development impacts DRG response to sensory stimulus (DMSO vs Jedi2: p=0.0427 unpaired t test) (DMSO n=12 animals, GsMTx4 n=11 animals, Jedi2 n=13 animals) (Fig 4Q). The noted nonsignificant decrease to the average length of shivering following GsMTx4 may be due in part from unidentified pathways targeted with this broad mechanosensitive ion channel antagonist. Overall these results support the hypothesis that impacting DRG development via Piezo1-mediated isolated Ca^2+^ transients results in a functional consequence to the animal’s physiology.

## Discussion

Activity of neural cells during development is well documented. We define this activity in satellite glia as significant changes in Ca^2+^ transients, which is a separate and different process from known neuronal firing activity. This activity can be broadly categorized into evoked and spontaneous Ca^2+^ transients. In glia, spontaneous Ca^2+^ transients are further divided into subtypes characterized as whole cell and microdomain Ca^2+^ transients. However, the developmental, molecular and functional features of these glial Ca^2+^ transients merits more investigation. Here, we demonstrate that satellite glia in the DRG exhibit distinct subtypes of spontaneous Ca^2+^ transients during early developmental times. We further reveal that distinct subtypes of Ca^2+^ transients are sensitive to manipulation of Piezo1. The functional consequence of disrupting such Piezo1-sensitive events is supported by data that shows cell abundance and sensory behavior is impacted by Piezo1-manipulations. Overall, we reveal developmental, molecular and functional characteristics of glial Ca^2+^ transients in the DRG.

Despite clear roles of neural activity in development and homeostasis of the nervous system in the animal, it is unclear when and which cells in the DRG show spontaneous Ca^2+^ transients. It is well appreciated that cultured DRG neurons exhibit spontaneous activity[54,55]. Ca^2+^ reporters have also demonstrated that satellite glia demonstrate Ca^2+^ transients in culture[56,57]. These cultured satellite glia also exhibit synchronized Ca^2+^ transients. Recent work has also shown Ca^2+^ transients in the vertebrate DRG neurons *in vivo*[58,59]. However, such work was restricted to mature animals. Our work reveals that both DRG neurons and glia in the animal are active during the earliest stages of DRG construction. Even on the first day of genesis, DRG cells demonstrate Ca^2+^ transients. We also identify the molecular mechanisms that mediate some of these Ca^2+^ transients. What remains unknown is how these Ca^2+^ transients change as the animal approaches adulthood or in neuropathologies. These are important topics to study because we know altered activity of both glia and neurons has been implicated in neuropathologies. Further, in addition to the spontaneous Ca^2+^ transients we focus on, neurons in the DRG also are evoked by specific stimuli. How glia respond to evoked stimulation in the animal is almost entirely unknown.

By imaging GCaMP6s in *sox10^+^* cells and probing gap junction components, we reveal that synchronized cellular Ca^2+^ transients indicative of glial networks form within 3 days of DRG genesis. We know that glial networks are essential in the central nervous system for circuit formation, neuronal health, and signal transduction. However, glial networks in the peripheral nervous system are less understood. Because of this, whether glial networks exist in developing DRG was not known. Our work identifies that DRG satellite glia are not synchronized initially, but by 4 days of DRG construction, become synchronized. One interesting aspect of this increased synchronization is that it occurs while the population is simultaneously expanding via cell divisions. Further investigation in additional PNS populations will provide an understanding if this is a unique process to the DRG, or if it is found in additional areas of the peripheral nervous system. It will also be important to probe the plasticity of the synchronized glial network and how it could be altered.

While most currently published research has focused on identifying molecules involved in synchronous Ca^2+^ transients of cells that make up neural circuits, we have identified a mechanism that contributes specifically to what we define as isolated Ca^2+^ transients within the DRG. We found that mechanobiology via Piezo1 contributes to an increase in isolated Ca^2+^ transients without altering simultaneous Ca^2+^ transients. Our findings provide insights into the importance of understanding mechanical forces on the cellular level during development. It is possible that these mechanical forces provide insight to satellite glia regarding whether proliferation is needed. For example, if the mechanical forces acting on a satellite glia are high, this may cause signaling via Piezo1 to halt or promote proliferation[29,35,60]. Alternatively, the ability to sense larger mechanical forces on the level of tissues may be important for DRG expansion. If this were true, an increase in mechanical forces may signal to the DRG that there is no room for further proliferation. One potential signaling cascade that may be involved in YAP/Taz. YAP/Taz is a well-known controller of cellular proliferation and has also been shown to modulate DRG development[13]. So, it is possible that over activating Piezo1 is altering localization of Yap/Taz[61]. Another area of research that would impact these ideas is the utility of Ca^2+^ microdomains observed in the DRG. If these microdomains are indicative of mechanical forces on the subcellular level, we may hypothesize either an increase or decrease in the amount of these microdomains which would further inform proliferative decisions in DRG satellite glia. Further investigation into the mechanical forces acting on the microenvironment of the DRG needs to be completed.

Our findings highlight the importance of understanding the role of mechanical signals in PNS development and the function of distinct Ca^2+^ transients in that process.

## Limitations of findings

We found a transition from an asynchronous population to a synchronous population during early DRG development. We attribute this transition to an increase in correlation coefficients between cells present in the DRG, which we hypothesize is a result from increased gap junction formation. But, this transition may also be partially explained by other processes. This work only investigates Ca^2+^ transients in the first 3 days of DRG genesis, so whether the observed synchrony remains through later stages of development is unknown and merits further study. We currently hypothesize that the decrease in cell proliferation is a result of Piezo1 overactivation. Whether Piezo1 controls proliferation only through isolated Ca^2+^ transients cannot be determined with our experiments. Our pharmacological manipulations also do not distinguish between cell-autonomous and non-autonomous roles of Piezo1 in satellite glia. Because of this, we cannot rule out the possibility that some observed phenotypes are a result of non-autonomous signaling. However, the *piezo1* expression in DRG *sox10^+^* cells and response to Piezo1 agonists, as well as our genetic manipulations, suggests a cell-autonomous role. But it remains a possibility that over activating Piezo1 could result in a change in a cell’s ability to communicate unidentified signals to surrounding cells. If this is the case, then non-autonomous roles of Piezo1 could contribute to the reduction in cell abundance. Lastly, we identify satellite glia as *sox10^+^ neurod^−^* cells residing in the DRG with an ensheathing morphology. It is likely that a subset of progenitor cells exist in the DRG during this developmental time period and potentially into adulthood.[37] If this is the case, our findings would suggest that altering Piezo1-mediated Ca^2+^ transients in both satellite glia and DRG progenitors likely impact DRG development.

## ACKNOWLEDGEMENTS

We thank members of the Wingert, Patzke and Smith labs for helpful discussions. Thank you to David Lyons for sharing GCaMP6s transgenic zebrafish. We also thank 3i for fielding imaging related questions and Deborah Bang, Brittany Gervais and others for zebrafish housing and upkeep. This work was supported by The Garibaldi Family Endowment for Excellence in Adult Stem Cell Research (JPB), The Hiller Family and Garibaldi Family Fellowships in Stem Cell and Regenerative Biology (JPB), Michael and Elizabeth Gallagher Family (CJS), The University of Notre Dame (CJS), the SMART foundation (CJS), and the NIH (DP2NS117177)(CJS). The funders had no role in study design, data collection, analysis, decision to publish or preparation of the manuscript.

## AUTHOR CONTRIBUTIONS STATEMENT

JPB performed all the analysis and experimentation, wrote the paper and conceived the study. CJS wrote and edited the manuscript and supervised and funded the project.

## COMPETING INTERESTS STATEMENT

The authors declare no competing interests.

## MATERIALS AND METHODS

Experimental Model and subject details

Experimental procedures adhered to the NIH guide for the care and use of laboratory animals. All experiments were approved by the University of Notre Dame Institutional Animal Care and Use Committee (IACUC) (protocol 19-08-5464) which is guided by the United States Department of Argriculture, the Animal Welfare Act (USA) and the Assessment and Accreditation of Laboratory Animal Care International.

Animal Specimens. Danio rerio (zebrafish) were utilized in this study. The following stable strains were used: AB, *Tg(sox10:gal4+myl7:gfp)*[62], *Tg(uas:GCaMP6s)*[63], *Tg(neurod:gal4+myl7:gfp)*[64]*, Tg(sox10:meGFP)*[65], *Tg(neurod:tagRFP)*[66]*, Tg(ngn1:GFP)*[67]. All embryos were produced through pairwise matings and grown in 28°C in constant darkness. At 24 hpf, zebrafish were exposed to PTU (0.0003%) to reduce pigmentation for intravital imaging. Age of animals was determined by hour post fertilization and stages of development[68].

### Experimental Procedures

#### *In vivo* overnight imaging

Animals were anesthetized using veterinary grade 3-aminobenzoic acid ester (Tricaine) for mounting purposes only. Animals were then mounted laterally on their right side in glass-bottomed 35 mm petri dishes[38] and covered in 0.8% low melt agarose. For overnight time lapse imaging, a mixture of egg water and tricaine was added to the dish. Images were acquired on spinning disk confocal microscopes custom built by 3i technology (Denver, CO) that contains: Zeiss Axio Observer Z1 Advanced Mariana Microscope, X-cite 120LED White Light LED System, filter cubes for GFP and mRFP, a motorized X,Y stage, piezo Z stage, 20X Air (0.50 NA), 63X (1.15NA), 40X (1.1NA) objectives, CSU-W1 T2 Spinning Disk Confocal Head (50 μm) with 1X camera adapter, and an iXon3 1Kx1K EMCCD camera or Prime 95B back illuminated CMOS camera, dichroic mirrors for 446, 515, 561, 405, 488, 561,640 excitation, laser stack with 405 nm, 445 nm, 488 nm, 561 nm and 637 nm. Overnight time-lapse images were collected every 5 min for 24 hours capturing a 40μm z stack. Adobe Illustrator and ImageJ were used to process images. Only brightness and contrast were adjusted and enhanced for images represented in this study.

#### *in vivo* calcium imaging

Animals were anesthetized using veterinary grade 3-aminobenzoic acid ester (Tricaine) for mounting purposes only. Animals were then mounted laterally on their right side in glass-bottomed 35 mm petri dishes[38] and covered in 0.8% low melt agarose. For Ca^2+^ imaging, egg water was added to the dish with no Tricaine. Images were acquired on a spinning disk confocal microscope custom built by 3i technology (Denver, CO) microscopes. Ca^2+^ time-lapse imaging consisted of image collection every 15 seconds for 1 hour capturing a 40μM z stack. For imaging of Ca^2+^ microdomains, images were either taken every 5s for 10 minutes capturing a 20μm z stack (Fig 1) or taken every 200ms in a single plane (Fig 3). Adobe Illustrator and ImageJ were used to process images. Only brightness and contrast were adjusted and enhanced for images represented in this study.

#### Pharmacological treatments

For this study, we used chemical treatments of HMR1556 20μM (Sigma-Aldrich), Thapsigargin 10μM (Sigma-Aldrich), Apyrase 10U (Sigma-Aldirch), Carbenoxolone 100μM (Tocris), GsMTx4 1μM (Tocris), Yoda1 100μM (Tocris), and Jedi2 40μM (Tocris). Concrentations were based on previous work and from testing a variety of concentrations of Piezo1 agonists (Sup Fig 6). HMR1556 was stored at 10mM concentration at −20°C. Thapsigargin was stored at 10mM concentration at −20°C. Apyrase was stored at 100U in −20°C. GsMTx4 was stored at 1mM at −20°C. Yoda1 was stored at 10mM at 4°C in 100% DMSO. Jedi2 was stored at 10mM at 4°C. All treatments were done in 2% DMSO. Control groups were treated with 2% DMSO throughout.

#### Consecutive Treatments of Pharmacological treatments

For consecutive days of treatment with pharmacological treatments animals were bathed in a mixture of either 40μM Jedi2, 1μM GsMTx4, or 100μM Yoda1 in egg water with 2% DMSO for 30 minutes each day. For Fig 4A-4F only the 40μM Jedi2 and 1μM GsMTx4 mixtures or a 2% DMSO egg water control were used. These treatments occurred at 2dpf and 3dpf. For Fig 4G-4J mixtures of 40μM Jedi2, 1μM GsMTx4, and 100μM Yoda1 mixtures or a 2% DMSO egg water control were used. These treatments occurred at 2dpf, 3dpf, and 4dpf. For Fig 4K-4L animals were treated with either 40μM Jedi2 or 2% DMSO egg water at 2 and 3dpf. For Fig 4M animals were treated with 40μM Jedi2, 1μM GsMTx4, or 2% DMSO egg water at 2dpf, 3dpf, and 4dpf.

#### Whole-mount immunohistochemistry

The primary antibody used to identify formation of gap junctions was Cxn43 (1:500; Cell Signaling Technology) (Fig 1O). The primary antibody used to identify cells expressing *Tg(sox10:meGFP)* was GFP (1:500; Aves) (Fig 2 H) following the listed RNAscope protocol. The primary antibody used to identify non-neuronal cells present in the DRG was Sox10 (1:1000, Sarah Kucenas Lab) (Fig 4H). The primary antibody used to identify neurons present in the DRG was HuCD (1:500, Thermo Fisher) (Fig 4L) The secondary antibody used in in Fig2H and Fig4H was Alexa Fluor 488 (1:500; Invitrogen). The secondary antibody used in Fig4L was Alexa Fluor 561 (1:500; Invitrogen). Animals were fixed in 4% PFA in PBSt (PBS, 0.1% TritonX-100) at 2dpf (Fig 2G) or 5dpf (Fig 4H, 4L). Animals were then washed for 5 minutes in a series of PBSt, DWt (dH2O, 0.1% TritonX-100), and acetone. Following these washes, animals were placed in acetone at −20°C for 10 minutes. This was then followed by 3 washes of PBSt for 5 minutes each. Animals were then placed in 5% goat serum in PBSt for a 1 hour incubation period. Animals were then incubated in 5% goat serum in PBSt with primary antibody for 1 hour at room temperature followed by an overnight incubation at 4°C. This was followed by 3 consecutive 30 minutes washes of PBSt and one 1 hour wash in PBSt. Animals were then placed in 5% goat serum in PBSt with the secondary antibody for 1 hour at room temperature followed by an overnight incubation at 4°C. This was then followed by 3 consecutive 1 hour washes of PBSt. Animals were then stored in 50% glycerol 50% PBS at 4°C until imaging.

#### Whole-mount RNAscope

Animals expressing *Tg(sox10:meGFP)* were fixed at 2dpf with 4%PFA in PBS for 30 minutes. Following fixation animals were placed in new eppendorf tubes and washed with 25%, 50%, 100% methanol for 10 minutes each. Animals were then kept in 100% methanol at −20°C overnight. This was followed by a 5 minute wash of 50% methanol in PBStw (PBS, 0.1% Tween-20) and an additional 5 minute wash of 25% methanol in PBStw. Liquid was removed from the eppendorf tubes and the animals were air dried for 30 minutes. This was followed by two 5 minute washes with PBStw. Animals were permeabilized with 10μg/mL proteinase K in PBStw at room temperature for 6 minutes. This was then followed by 4 consecutive 10 minute washes of PBStw. Following removal of PBStw, 2 drops of ACD probes targeting *piezo1* were then added to the sample, which was then incubated at 40°C for 15 hours (1:50, 80uL, C1, ACD). In the case of positive and negative controls that were used, Probe-Dr-polr2 was used as a ubiquitous positive control and Probe-Dr-dapB was used as a bacterial negative control. After this incubation period animals were then washed with SSCtw (5X saline-sodium citrate buffer, 0.1% Tween-20) for 10 minutes at room temperature twice. An additional fixation was then done in 4% PFA in PBS at room temperature for 10 minutes. Animals were again transferred to a new eppendorf tube and washed 3 times in SSCtw. Animals were then incubated in a series of 2 drops Amp1 at 40°C for 30 minutes, 2 drops Amp2 at 40°C for 30 minutes, 2 drops Amp3 at 40°C for 15 minutes, 2 drops HRP-C1 at 40°C for 30 minutes, opal fluorophore 650 (1:500) in PBStw at 40°C for 30 minutes, and 2 drops of Multiplex FLv2 HRP blocker at 40°C for 30 minutes. Between each of these incubation periods animals were washed twice with SSCtw. Following this protocol, animals were then processed following the Whole-mount immunohistochemistry protocol to target GFP.

#### Whole-mount HCR-FISH

Animals expressing *Tg(sox10:megfp)* were fixed at 3dpf in 4% PFA in PBS for 24 hours at 4°C. Fixed larvae were then washed in PBS 3 times for 5 minutes each. To dehydrate and permeabilize the tissue samples were then washed in a series of 100% methanol 4 times for 10 minutes each. Samples were then stored in 100% methanol at −20°C for 24 hours. To rehydrate samples a series of methanol/PBStw washes were utilized. Samples were washed for 5 minutes in 75% methanol in PBStw, 50% methanol in PBStw, 25% methanol in PBStw, and 100% PBStw. Samples were then treated with 10μg/mL proteinase K for 30 minutes. Samples were then washed in PBStw twice for 5 minutes. A postfix was done on samples in 4%PFA for an additional 20 minutes. Samples were then washed 5 times for 5 minutes each with PBStw. Samples were then washed in probe hybridization buffer (Molecular Instruments) for 30 minutes at 37°C. Samples were then incubated at 37°C overnight in probe hybridization buffer containing HCR-FISH probes targeting *piezo1* (Molecular Instruments). Samples were washed following incubation with probe wash buffer (Molecular Instruments) at 37°C 4 times for 15 minutes each. Samples were then washed twice with SSCtw for 5 minutes each. Following these washes samples were then incubated in amplification buffer (Molecular Instruments) for 30 minutes. Following this incubation samples were then incubated over night in amplification buffer containing hairpin b1h1 and hairpin b1h2 (647) overnight. Samples were then washed 5 times in PBStw for 20 minutes each. Following this protocol samples then underwent the Whole-mount immunohistochemistry protocol targeting GFP.

#### Animal behavior in cold stimulus

Animals were treated at 2, 3, and 4 dpf with 1μM GsMTx4 in 2% DMSO, 40μM Jedi2 in 2% DMSO, or 2% DMSO in egg water for 30 minutes each day. Aside from this treatment protocol, animals were raised under normal procedures in 28°C egg water. Each animal was then taken at 5dpf and placed in 4°C egg water for 30 seconds. Video recordings were recorded with 40ms exposure to bright field white light. The initial 3 seconds of the movies were not quantified to allow the animal to be placed into the stimulus and for any adjustments to occur. The duration of the shivering was quantified per second starting from the initial shivering phenotype to the cold water stimulus until there was at least 1 second of no shivering.

#### Mechanical compression assay

To apply mechanical force to the DRG, we dorsally mounted animals in 0.8% low melt agarose and placed them on the stage of a spinning disk confocal. On either side of the stage two vertical stainless steel rods were mounted onto the air table. A horizontal rod was mounted above the stage with a micromanipulator attached. A glass needle filled with dextran was mounted in the micromanipulator above the animals with the needle pointing toward the animal. Prior to bringing into contact with the animal, the needle was calibrated in the X and Y positions in relation to the middle of the imaging window. The needle was slowly brought into contact with the skin of the animals and was then tapped to apply pressure to the DRG and surrounding tissue. To understand if DRG were active in response to the mechanical forces, we quantified GCaMP6s transients as previously described. As a proxy to understand how much force was needed to activate DRG satellite glia, we quantified the response of DRG to a gradient of force measured by the distance (μm) that the DRG was displaced with animal bending in the XY direction.

#### Microinjections

In order to label cell membranes with GCaMP6s, animals expressing *Tg(sox10:gal4+myl7:GFP)* were injected with *uas:GCaMP6s-caax* at the single cell stage. In order for cell-specific CRISPR/Cas9, *uas:cas9mkate-u6:piezo1gRNA* or *uas:cas9mkate-u6:emptygRNA* were injected into animals expressing *Tg(sox10:gal4); Tg(uas:GCaMP6s)*. Injection mixes consisted of either 12 ng/μL *uas:GCaMP6s-caax* or *uas:cas9mkate-u6:piezo1gRNA* or *uas:cas9mkate-u6:emptygRNA* 25ng/μL *tol2* (transposase), and phenol red (visualization). Additional injection mixes consisted of 12/ng/μL *uas:Chr2-tdTomato*, 25ng/μL *tol1* (transposase), and phenol red. These injections were used for optogenetic manipulations. Animals were screened for expression of appropriate expression of incorporated transgenes and used in experiments.

#### Optogenetic manipulation

For genetic manipulations utilizing *channelrhodopsin*, we injected a *uas:Chr2-tdTomato* (Addgene 124237) into animals expressing *Tg(sox10:gal4); Tg(uas:GCaMP6s)*.[51] Animals were screened for *uas:Chr2-tdTomato* and either exposed to 488nm light for 30 minutes each day from 2-4dpf or not exposed to 488nm light each day from 2-4dpf.

#### Cell-specific CRISPR Cas9

In order to target *piezo1* specifically in our cells of interest, we utilized the *gal4/uas* system. A plasmid was constructed utilizing the Gateway LR Clonease II Plus system (ThermoFisher). To create *uas:cas9mkate-u6:piezo1gRNA*, we recombined a *p5E-UAS* (BseRI cleavage site removed), *pME-cas9mkate* (Addgene 109547), and *p3E-pA* constructs into a *pDestTol2CG2* containing a *U6* promoter sequence flanked with BseRI cleavage sites.[50,64] Following recombination the resulting plasmid, *uas:cas9mkate-u6:emptyGRNA* was digested with BseRI and annealed primers of *piezo1* gRNA (*piezo1* forward primer: GCACCCTGAGGATCTTCCAGGT, *piezo1* reverse primer: CTGGAAGATCCTCAGGGTGTGA) were ligated into the vector to create *uas:cas9mkate-u6:piezo1gRNA*. A control plasmid *uas:cas9mkate-u6:emptygRNA* with an empty gRNA cassette was used.

### Quantifications and Statistical Tests

#### Quantification of GCaMP6s transients

Before quantifying changes in GCaMP6s intensity, we corrected for motion drift by utilizing the Template Matching plugin in ImageJ. We then traced individual cells in the DRG to create Regions of Interest (ROI). The integrated density of fluorescence was quantified at every time point for each ROI. We then quantified the z score for each timepoint for each ROI. Timepoints with a z score of 2.58 or greater were considered active timepoints. A single activity event was identified by timepoints with a z score of 2.58 or greater. If consecutive timepoints were a z score of 2.58 or greater this was still considered one activity event. This process was used to create activity profiles for each ROI in a given DRG. The average number of active timepoints was calculated and compared to controls via t tests to determine significance.

#### Quantification of GCaMP6s transients in pilot screen

The number of GCaMP6s transients in the pilot screen were quantified in ImageJ. The number of visual changes in GCaMP6s intensity were quantified for each movie. These quantifications were done per whole DRG identified in the imaging window.

#### Generation of line graphs

Line graphs were generated using the data obtained from the quantification of GCaMP6s transients. Each cell in the DRG had GCaMP6s transients quantified. The X axis of the line graph is the 1 hr period and the Y axis is the Z score. Line graphs were generated in Prism.

#### Generation of Heatmaps

Heatmaps were generated in Prism. Each row of the heatmap corresponds to an individual cell's GCaMP6s transience during the 1hr of imaging. Blue colors indicate low z scores and yellow colors indicate high z scores (>2.58).

#### Quantification of Correlation Coefficients

Activity profiles from individual cells were converted to a binary system. Active points where the z score was greater than 2.58 were listed as 1 and inactive time points where the z score was less than 2.58 were listed as 0. These binary activity profiles were then used to quantify the correlation between individual ROIs found in the same DRG; we utilized the cor() function in R to complete this analysis. The results were then used to create a correlation table in R. These functions were part of the Hmisc package in R. Correlation coefficients greater than 0.5 were considered high correlation coefficients. The percent of high correlation coefficients were then quantified per cell and compared with controls via t tests to determine significance.

#### Quantification of IHC, RNAscope, HCR-FISH

All quantifications of IHC were completed in ImageJ. The number of Cxn43 puncta that coincided with *sox10:megfp* expression were quantified per DRG (Fig 1O,P). The number of DRG that contained *piezo1* within the stained GFP expressed was quantified (Fig 2H,I, Sup Fig 4). This was done by going through each individual optical slice to identify *piezo1* puncta within *sox10^+^* cells located in the DRG. The number of Sox10^+^ cells were quantified per DRG (Fig 4H,I). The number of HuCD^+^ cells were quantified per DRG (Fig M,N).

#### Quantification of Mechanical Compression Assay

Response to mechanical compression was quantified in ImageJ. DRG were traced and the integrated density of fluorescence was quantified at each time point. To attempt quantifying only time points in the same z position, the beginning and end time points quantified were in the same z position that were not manually being changed using the microscope. Due to the nature of this experiment, tissue compression would still alter the z position of the DRG being quantified. To further account for this alteration, the change in integrated density of fluorescence was quantified by subtracting the initial time point from each time point being analyzed (*Δf* = *f* − *fi*). This value was then divided by the time point being analyzed (*Δf*/*f*). Large increases in this value following tissue compression were then scored as active responses to mechanical compression. Little to no change in this value following mechanical compression were scored as not active in response to tissue compression.

#### Quantification of satellite glia ensheathment

The percent of *sox10^+^* ensheathment around *neurod+* cells was quantified utilizing ImageJ tracings. These quantifications were taken from overnight timelapse imaging of animals expressing *Tg(sox10:gal4+myl7); Tg(uas:GCaMP6s); Tg(neurod:tagRFP)* at 3dpf (Sup Fig 2B). The percent of *sox10^+^* cells found in a DRG with a wrapping phenotype were quantified as ensheathing satellite glia. These quantifications were taken from *in vivo* calcium imaging of animals expressing *Tg(sox10:gal4+myl7); Tg(uas:GCaMP6s); Tg(neurod:tagRFP)* at 3dpf following treatment of 40μM Jedi2 (Sup Fig 2C).

#### Statistical analysis

Statistical analysis was completed with Prism. No statistical methods were used to predetermine sample sizes but sample sizes are similar to previous publications. Statistical tests were completed with biological replicates, not technical replicates. No data points were excluded from the analysis. Healthy animals were randomly selected for all experiments. Each experiment was repeated at least with similar results. All data collected and analyzed are presented in the study.

#### Software

Slidebook, Prism, ImageJ, R, and Adobe Illustrator were used to acquire, analyze and compile figures.

## DATA AVAILABILITY

All data collected for this study are included in the figures and supplementary material.

## SUPPLEMENTAL FIGURES

**Supplemental Figure 1:**
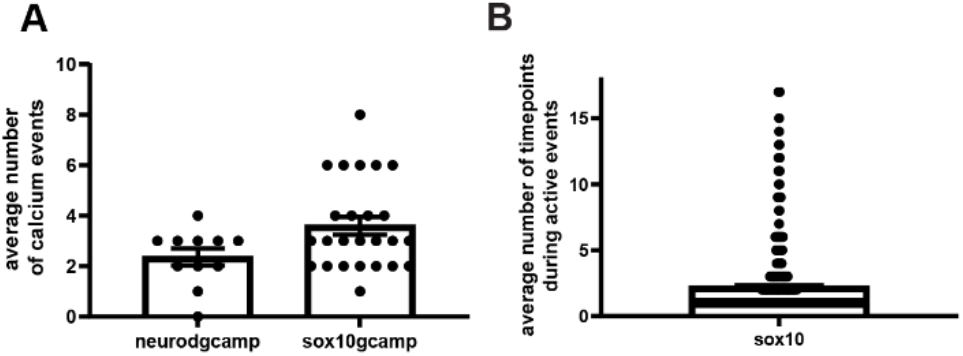
DRG cells are spontaneously active. A) Quantification of the average number of calcium events per *neurod^+^* cell or per *sox10^+^* cell in 3dpf animals expressing either *Tg(neurod:gal4+myl7); Tg(uas:GCaMP6s) or Tg(sox10:gal4+myl7); Tg(uas:GCaMP6s)*. (*neurod*: 4 animals, 9 DRG, 11 cells, *sox10*: 5 animals, 14 DRG, 25 cells) B) Quantification of the average number of timepoints during active events per *sox10^+^* cells in animals expressing *Tg(sox10:gal4+myl7); Tg(uas:GCaMP6s)*. (5 animals, 20 DRG, 97 cells, 412 calcium transient events)

**Supplemental Figure 2:**
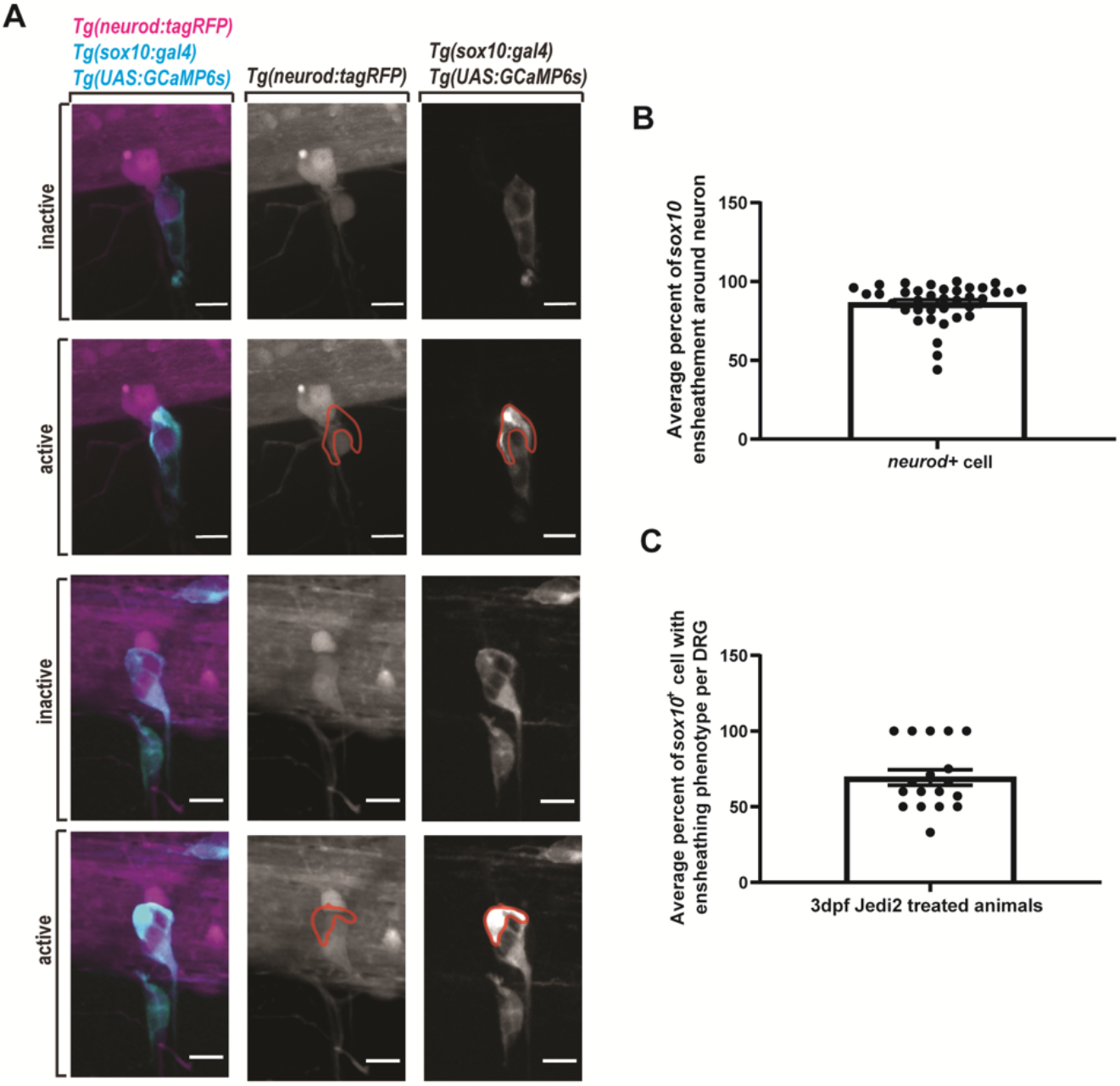
DRG contain *sox10^+^* satellite glia with ensheathing morphologies. A) confocal z-projections of DRG in 3dpf animals expressing *Tg(sox10:gal4+myl7); Tg(uas:GCaMP6s); Tg(neurod:tagRFP)*. Magenta displays *neurod^+^* neurons. Cyan displays *sox10^+^* satellite glia. Red tracing indicates the morphology of a satellite glia during a Ca^2+^ transient event. B) Quantification of the average percent of *sox10^+^* satellite glia ensheathment around a *neurod^+^* neuron. (5 animals, 14 DRG, 37 cells) C) Quantification of the average number of *sox10^+^* cells with an ensheathing morphology following 40μM Jedi2 treatment. (7 animals, 18 DRG, 74 cells) Scale bar is 10μm (A).

**Supplemental Figure 3:**
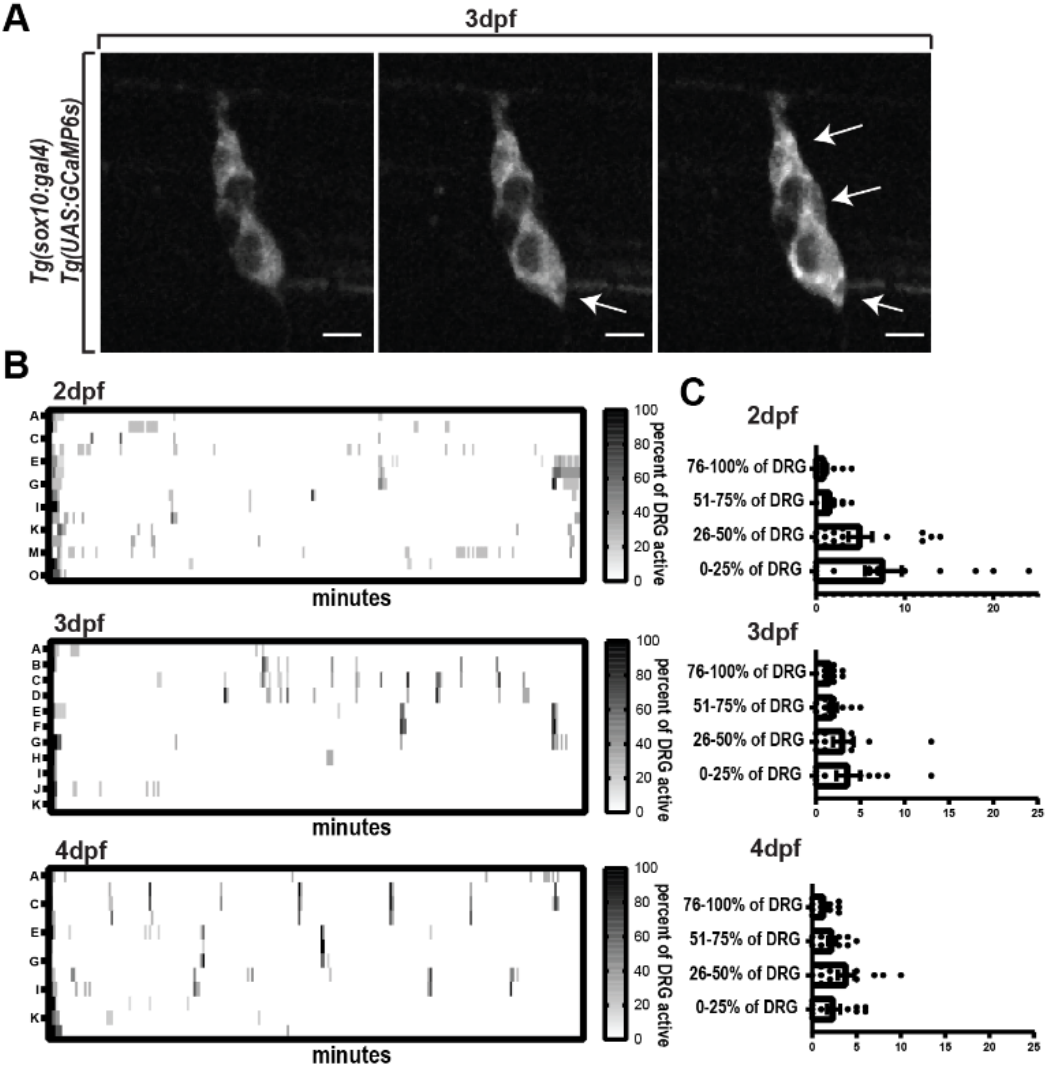
Percent of active DRG cells increase during development. A) Confocal z-projection of DRG in 3dpf animals expressing *Tg(sox10:gal4+myl7); Tg(uas:GCaMP6s).* Arrows note active cells and demonstrate different percentages of active DRG. B) Heatmaps of the percent of cells in the DRG active during a 1-hour period at 2, 3, and 4dpf. Darker gradient indicates a higher percent of cells active. (2dpf: 6 animals, 10 DRG, 46 cells, 3dpf: 4 animals, 6 DRG, 27 cells, 4dpf: 4 animals, 7 DRG, 34 cells) C) Quantification of the number of active events with 0-25%, 26-50%, 51-75%, or 76-100% of DRG cells active at the same time point at 2, 3, and 4dpf. . (2dpf: 6 animals, 10 DRG, 46 cells, 3dpf: 4 animals, 6 DRG, 27 cells, 4dpf: 4 animals, 7 DRG, 34 cells) Scale bar is 10μm (A)

**Supplemental Figure 4:**
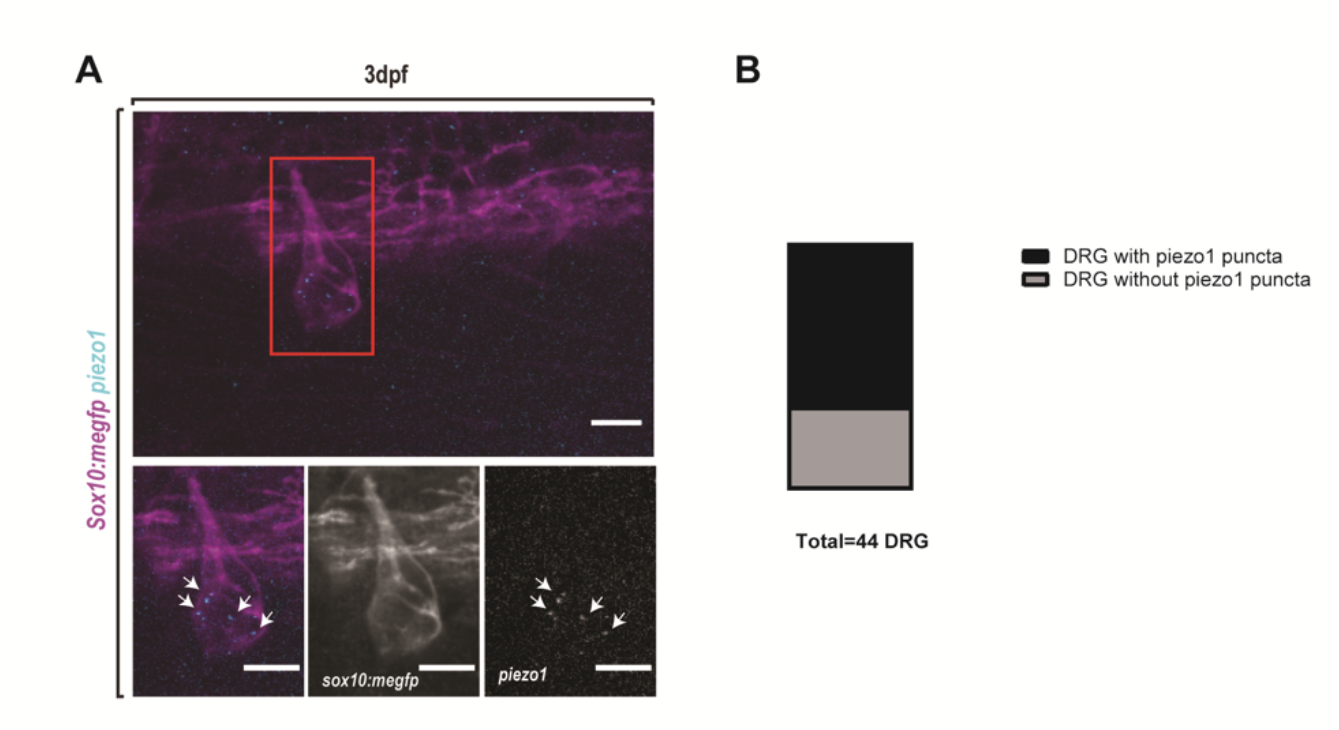
Whole-mount HCR-FISH targeting *piezo1* in 3dpf DRG. A) Confocal images of HCR-FISH-*piezo1* and Immunohistochemistry-GFP in 3dpf *Tg(sox10:meGFP)* animals. GFP is shown in magenta and *piezo1* is shown in cyan. Arrows indicate *piezo1* puncta. B) Quantification of DRG at 3 dpf with *piezo1* puncta and without *piezo1* puncta. (13 animals, 44 DRG) Scale bar is 10μm (A).

**Supplemental Figure 5:**
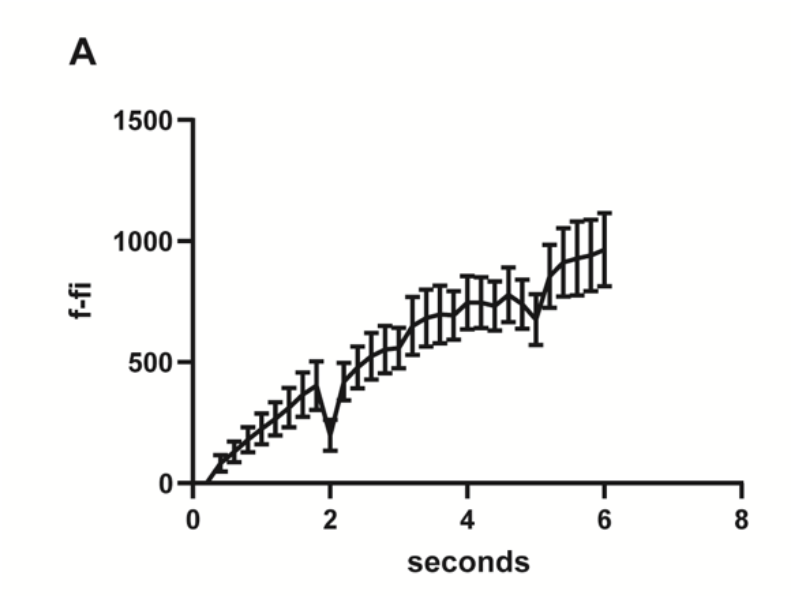
Change in fluorescence following cochr2 activation via 488nm light. A) Quantification of the average change in integral density of fluorescence over time (seconds). (7 animals, 7 DRG, 7 cells)

**Supplemental Figure 6:**
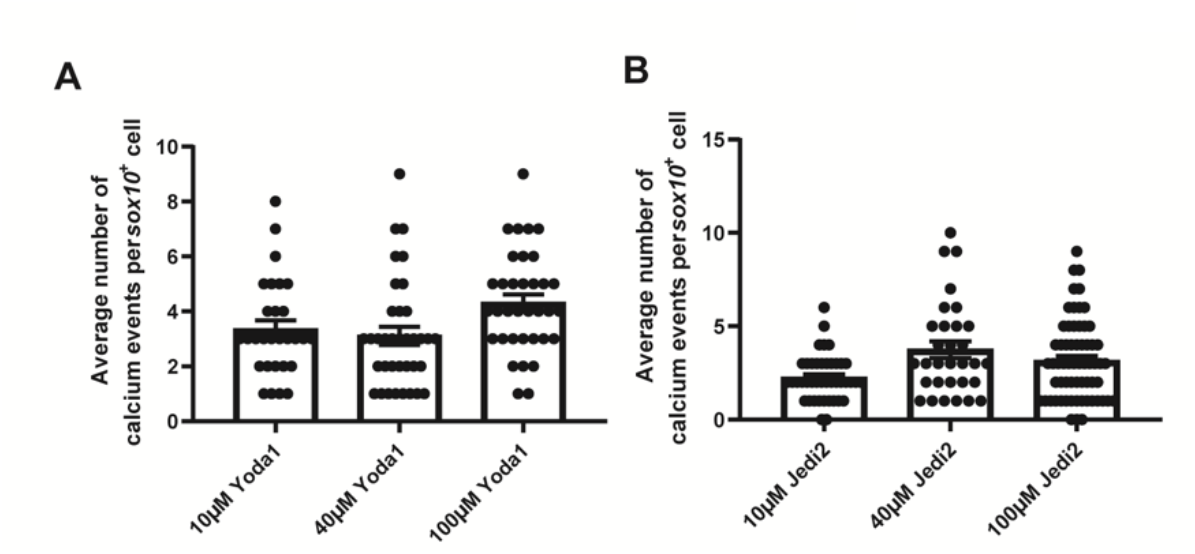
Increasing concentrations of Piezo1 agonists. A) Quantification of the average number of Ca^2+^ transient events in *sox10^+^* cells of 3dpf animals expressing *Tg(sox10:gal4+myl7); Tg(uas:GCaMP6s); Tg(neurod:tagRFP)* treated with 10μM, 40μM, or 100μM Yoda1 in 2%DMSO for 30 minutes prior to imaging. (10μM: 2 animals, 7 DRG, 29 cells, 40μM: 3 animals, 9 DRG, 36 cells, 100μM: 2 animals, 5 DRG, 35 cells) B) Quantification of the average number of Ca^2+^ transient events in *sox10^+^* cells of 3dpf animals expressing *Tg(sox10:gal4+myl7); Tg(uas:GCaMP6s); Tg(neurod:tagRFP)* treated with 10μM, 40μM, or 100μM Jedi2 in 2%DMSO for 30 minutes prior to imaging. (10μM: 3 animals, 9 DRG, 42 cells, 40μM: 3 animals, 8 DRG, 31 cells, 100μM: 5 animals, 14 DRG, 63 cells)

**Supplemental Table 1. Statistics for Study.** Summary of statistical information for each figure panel.

## Notes

### Competing Interest Statement

The authors have declared no competing interest.

### Summary of Updates

Updated version includes additional experiments.

## REFERENCES

1. Triplett MA, Avitan L, Goodhill GJ. Emergence of spontaneous assembly activity in developing neural networks without afferent input. PLoS Comput Biol. 2018;14: 1–22. doi:10.1371/journal.pcbi.1006421

2. Pietri T, Romano SA, Pérez-Schuster V, Boulanger-Weill J, Candat V, Sumbre G. The Emergence of the Spatial Structure of Tectal Spontaneous Activity Is Independent of Visual Inputs. Cell Rep. 2017;19: 939–948. doi:10.1016/j.celrep.2017.04.015

3. Avitan L, Pujic Z, Mo J, Amor R, Scott EK, Goodhill GJ, et al. Spontaneous Activity in the Zebrafish Tectum Reorganizes over Development and Is Influenced by Article Spontaneous Activity in the Zebrafish Tectum Reorganizes over Development and Is Influenced by Visual Experience. 2017; 2407–2419. doi:10.1016/j.cub.2017.06.056

4. Ma Z, Stork T, Bergles DE, Freeman MR. Neuromodulators signal through astrocytes to alter neural circuit activity and behaviour. Nature. 2016;539: 428–432. doi:10.1038/nature20145

5. Sasaki T, Ishikawa T, Abe R, Nakayama R, Asada A, Matsuki N, et al. Astrocyte calcium signalling orchestrates neuronal synchronization in organotypic hippocampal slices. J Physiol. 2014;592: 2771–2783. doi:10.1113/jphysiol.2014.272864

6. Stobart JL, Ferrari KD, Barrett MJP, Stobart MJ, Looser ZJ, Saab AS, et al. Long-term in vivo calcium imaging of astrocytes reveals distinct cellular compartment responses to sensory stimulation. Cereb Cortex. 2018;28: 184–198. doi:10.1093/cercor/bhw366

7. Kuijlaars J, Oyelami T, Diels A, Rohrbacher J, Versweyveld S, Meneghello G, et al. Sustained synchronized neuronal network activity in a human astrocyte co-culture system. Sci Rep. 2016;6: 1–14. doi:10.1038/srep36529

8. Ma Z, Freeman MR. Trpml-mediated astrocyte microdomain ca2+ transients regulate astrocyte-tracheal interactions. Elife. 2020;9: 1–18. doi:10.7554/ELIFE.58952

9. Rui Y, Pollitt SL, Myers KR, Feng Y, Zheng JQ. Spontaneous local calcium transients regulate oligodendrocyte development in culture through store-operated ca2+ entry and release. eNeuro. 2020;7: 1–16. doi:10.1523/ENEURO.0347-19.2020

10. Baraban M, Koudelka S, Lyons DA. Ca 2+ activity signatures of myelin sheath formation and growth in vivo. Nat Neurosci. 2018;21: 19–25. doi:10.1038/s41593-017-0040-x

11. Hösli L, Binini N, Ferrari KD, Thieren L, Looser ZJ, Zuend M, et al. Decoupling astrocytes in adult mice impairs synaptic plasticity and spatial learning. Cell Rep. 2022;38. doi:10.1016/j.celrep.2022.110484

12. Weiss S, Clamon LC, Manoim JE, Ormerod KG, Parnas M, Littleton JT. Glial ER and GAP junction mediated Ca2+ waves are crucial to maintain normal brain excitability. Glia. 2022;70: 123–144. doi:10.1002/glia.24092

13. Serinagaoglu Y, Paré J, Giovannini M, Cao X. Nf2-Yap signaling controls the expansion of DRG progenitors and glia during DRG development. Dev Biol. 2015;398: 97–109. doi:10.1016/j.ydbio.2014.11.017

14. Coronas V, Terrié E, Déliot N, Arnault P, Constantin B. Calcium Channels in Adult Brain Neural Stem Cells and in Glioblastoma Stem Cells. Front Cell Neurosci. 2020;14: 1–22. doi:10.3389/fncel.2020.600018

15. Venkatesh HS, Morishita W, Geraghty AC, Silverbush D, Gillespie SM, Arzt M, et al. Electrical and synaptic integration of glioma into neural circuits. Nature. 2019;573: 539–545. doi:10.1038/s41586-019-1563-y

16. Chen X, Wanggou S, Bodalia A, Zhu M, Dong W, Fan JJ, et al. A Feedforward Mechanism Mediated by Mechanosensitive Ion Channel PIEZO1 and Tissue Mechanics Promotes Glioma Aggression. Neuron. 2018;100: 799–815.e7. doi:10.1016/j.neuron.2018.09.046

17. Armstrong DM. Glutamate Receptor Subtypes Mediate Excitatory of Dopamine Neurons in Midbrain Slices Synaptic Currents. 1991; 3–4.

18. Cheramy A, Leviel V, Glowinski J. Dendritic release of dopamine in the substantia nigra. 1981;289: 537–542.

19. Micu I, Jiang Q, Coderre E, Ridsdale A, Zhang L, Woulfe J, et al. NMDA receptors mediate calcium accumulation in myelin during chemical ischaemia. Nature. 2006;439: 988–992. doi:10.1038/nature04474

20. Káradóttir R, Cavelier P, Bergersen LH, Attwell D. NMDA receptors are expressed in oligodendrocytes and activated in ischaemia. Nature. 2005;438: 1162–1166. doi:10.1038/nature04302

21. Rose CR, Felix L, Zeug A, Dietrich D, Reiner A, Henneberger C, et al. Astroglial Glutamate Signaling and Uptake in the Hippocampus. 2018;10: 1–20. doi:10.3389/fnmol.2017.00451

22. Gonzalez A, Almeida A, Bolaños JP. Astrocyte NMDA receptors ’ activity sustains neuronal survival through a Cdk5 – Nrf2 pathway. 2015; 1877–1889. doi:10.1038/cdd.2015.49

23. Bowser DN, Khakh BS. ATP Excites Interneurons and Astrocytes to Increase Synaptic Inhibition in Neuronal Networks. 2004;24: 8606–8620. doi:10.1523/JNEUROSCI.2660-04.2004

24. Zhang J, Wang H, Ye C, Ge W, Chen Y, Jiang Z, et al. ATP Released by Astrocytes Mediates Heterosynaptic Suppression. 2003;40: 971–982.

25. Puerto A, Wandosell F, Garrido JJ. Neuronal and glial purinergic receptors functions in neuron development and brain disease. 2013;7: 1–15. doi:10.3389/fncel.2013.00197

26. Koser DE, Thompson AJ, Foster SK, Dwivedy A, Pillai EK, Sheridan GK, et al. Mechanosensing is critical for axon growth in the developing brain. Nat Neurosci. 2016;19: 1592–1598. doi:10.1038/nn.4394

27. Espinosa-Hoyos D, Burstein SR, Cha J, Jain T, Nijsure M, Jagielska A, et al. Mechanosensitivity of Human Oligodendrocytes. Front Cell Neurosci. 2020;14: 1–15. doi:10.3389/fncel.2020.00222

28. Song Y, Li D, Farrelly O, Miles L, Li F, Kim SE, et al. The Mechanosensitive Ion Channel Piezo Inhibits Axon Regeneration. Neuron. 2019;102: 373–389.e6. doi:10.1016/j.neuron.2019.01.050

29. Pathak MM, Nourse JL, Tran T, Hwe J, Arulmoli J, Le DTT, et al. Stretch-activated ion channel Piezo1 directs lineage choice in human neural stem cells. Proc Natl Acad Sci U S A. 2014;111: 16148–16153. doi:10.1073/pnas.1409802111

30. Hill RZ, Loud MC, Dubin AE, Peet B, Patapoutian A. PIEZO1 transduces mechanical itch in mice. Nature. 2022;607: 104–110. doi:10.1038/s41586-022-04860-5

31. Wang J, La JH, Hamill OP. PIEZO1 is selectively expressed in small diameter mouse DRG neurons distinct from neurons strongly expressing TRPV1. Front Mol Neurosci. 2019;12: 1–15. doi:10.3389/fnmol.2019.00178

32. Abdo H, Calvo-Enrique L, Lopez JM, Song J, Zhang MD, Usoskin D, et al. Specialized cutaneous schwann cells initiate pain sensation. Science (80-). 2019;365: 695–699. doi:10.1126/science.aax6452

33. Ackerman SD, Perez-Catalan NA, Freeman MR, Doe CQ. Astrocytes close a motor circuit critical period. Nature. 2021;592: 414–420. doi:10.1038/s41586-021-03441-2

34. Shigetomi E, Tong X, Kwan KY, Corey DP, Khakh BS. TRPA1 channels regulate astrocyte resting calcium and inhibitory synapse efficacy through GAT-3. Nat Neurosci. 2012;15: 70–80. doi:10.1038/nn.3000

35. Rammensee S, Kang MS, Georgiou K, Kumar S, Schaffer D V. Dynamics of Mechanosensitive Neural Stem Cell Differentiation. Stem Cells. 2017;35: 497–506. doi:10.1002/stem.2489

36. Marinval N, Chew SY. Mechanotransduction assays for neural regeneration strategies: A focus on glial cells. APL Bioeng. 2021;5. doi:10.1063/5.0037814

37. Maniglier M, Vidal M, Bachelin C, Deboux C, Chazot J, Garcia-Diaz B, et al. Satellite glia of the adult dorsal root ganglia harbor stem cells that yield glia under physiological conditions and neurons in response to injury. Stem Cell Reports. 2022;17: 2467–2483. doi:10.1016/j.stemcr.2022.10.002

38. Nichols EL, Green LA, Smith CJ. Ensheathing cells utilize dynamic tiling of neuronal somas in development and injury as early as neuronal differentiation. Neural Dev. 2018;13: 19. doi:10.1186/s13064-018-0115-8

39. Mazaud D, Capano A, Rouach N. The many ways astroglial connexins regulate neurotransmission and behavior. Glia. 2021;69: 2527–2545. doi:10.1002/glia.24040

40. Mu Y, Bennett D V., Rubinov M, Narayan S, Yang CT, Tanimoto M, et al. Glia Accumulate Evidence that Actions Are Futile and Suppress Unsuccessful Behavior. Cell. 2019;178: 27–43.e19. doi:10.1016/j.cell.2019.05.050

41. Hughes AN, Appel B. Oligodendrocytes express synaptic proteins that modulate myelin sheath formation. Nat Commun. 2019;10: 1–15. doi:10.1038/s41467-019-12059-y

42. Hanani M, Spray DC. Emerging importance of satellite glia in nervous system function and dysfunction. Nat Rev Neurosci. 2020;21: 485–498. doi:10.1038/s41583-020-0333-z

43. Huang T, Belzer V, Hanani M. Gap junctions in dorsal root ganglia : Possible contribution to visceral pain. Eur J Pain. 2010;14: 49.e1–49.e11. doi:10.1016/j.ejpain.2009.02.005

44. Huang LYM, Gu Y, Chen Y. Communication between neuronal somata and satellite glial cells in sensory ganglia. Glia. 2013;61: 1571–1581. doi:10.1002/glia.22541

45. Retamal MA, Riquelme MA, Stehberg J, Alcayaga J. Connexin43 hemichannels in satellite glial cells, can they influence sensory neuron activity? Front Mol Neurosci. 2017;10: 1–9. doi:10.3389/fnmol.2017.00374

46. Komiya H, Shimizu K, Ishii K, Kudo H, Okamura T, Kanno K, et al. Connexin 43 expression in satellite glial cells contributes to ectopic tooth-pulp pain. J Oral Sci. 2018;60: 493–499. doi:10.2334/josnusd.17-0452

47. Ohara PT, Vit JP, Bhargava A, Jasmin L. Evidence for a role of connexin 43 in trigeminal pain using RNA interference in vivo. J Neurophysiol. 2008;100: 3064–3073. doi:10.1152/jn.90722.2008

48. Syeda R, Xu J, Dubin AE, Coste B, Mathur J, Huynh T, et al. Chemical activation of the mechanotransduction channel Piezo1. Elife. 2015;4: 1–11. doi:10.7554/eLife.07369

49. Wang Y, Chi S, Guo H, Li G, Wang L, Zhao Q, et al. A lever-like transduction pathway for long-distance chemical- and mechano-gating of the mechanosensitive Piezo1 channel. Nat Commun. 2018;9. doi:10.1038/s41467-018-03570-9

50. Ablain J, Durand EM, Yang S, Zhou Y, Zon LI. A CRISPR/Cas9 vector system for tissue-specific gene disruption in zebrafish. Dev Cell. 2015;32: 756–764. doi:10.1016/j.devcel.2015.01.032

51. Antinucci P, Dumitrescu AS, Deleuze C, Morley HJ, Leung K, Hagley T, et al. A calibrated optogenetic toolbox of stable zebrafish opsin lines. Elife. 2020;9: 1–31. doi:10.7554/eLife.54937

52. Kikel-Coury NL, Green LA, Nichols EL, Zellmer AM, Pai S, Hedlund SA, et al. Pioneer axons utilize a Dcc signaling-mediated invasion brake to precisely complete their pathfinding odyssey. J Neurosci. 2021;41: 6617–6636. doi:10.1523/JNEUROSCI.0212-21.2021

53. Nichols EL, Smith CJ. Pioneer axons employ Cajal’s battering ram to enter the spinal cord. Nat Commun. 2019;10. doi:10.1038/s41467-019-08421-9

54. Chen Y, Huang LYM. A simple and fast method to image calcium activity of neurons from intact dorsal root ganglia using fluorescent chemical Ca2+ indicators. Mol Pain. 2017;13: 1–9. doi:10.1177/1744806917748051

55. Teichert RW, Smith NJ, Raghuraman S, Yoshikami D, Light AR, Olivera BM. Functional profiling of neurons through cellular neuropharmacology. Proc Natl Acad Sci U S A. 2012;109: 1388–1395. doi:10.1073/pnas.1118833109

56. Suadicani SO, Cherkas PS, Zuckerman J, Smith DN, Spray DC, Hanani M. Bidirectional calcium signaling between satellite glial cells and neurons in cultured mouse trigeminal ganglia. Neuron Glia Biol. 2010;6: 43–51. doi:10.1017/S1740925X09990408

57. Zhang X, Chen Y, Wang C, Huang LYM. Neuronal somatic ATP release triggers neuron-satellite glial cell communication in dorsal root ganglia. Proc Natl Acad Sci U S A. 2007;104: 9864–9869. doi:10.1073/pnas.0611048104

58. Chen C, Zhang J, Sun L, Zhang Y, Gan WB, Tang P, et al. Long-term imaging of dorsal root ganglia in awake behaving mice. Nat Commun. 2019;10: 1–11. doi:10.1038/s41467-019-11158-0

59. Chisholm KI, Khovanov N, Lopes DM, La Russa F, McMahon SB. Large scale in vivo recording of sensory neuron activity with GCaMP6. eNeuro. 2018;5: 1–14. doi:10.1523/ENEURO.0417-17.2018

60. Segel M, Neumann B, Hill MFE, Weber IP, Viscomi C, Zhao C, et al. Niche stiffness underlies the ageing of central nervous system progenitor cells. Nature. 2019;573: 130–134. doi:10.1038/s41586-019-1484-9

61. Dasgupta I, McCollum D. Control of cellular responses to mechanical cues through YAP/TAZ regulation. J Biol Chem. 2019;294: 17693–17706. doi:10.1074/jbc.REV119.007963

62. Hines JH, Ravanelli AM, Schwindt R, Scott EK, Appel B. Neuronal activity biases axon selection for myelination in vivo. Nat Neurosci. 2015;18: 683–689. doi:10.1038/nn.3992

63. Thiele TR, Donovan JC, Baier H. Descending Control of Swim Posture by a Midbrain Nucleus in Zebrafish. Neuron. 2014;83: 679–691. doi:10.1016/j.neuron.2014.04.018

64. Nichols EL, Smith CJ. Synaptic-like Vesicles Facilitate Pioneer Axon Invasion. Curr Biol. 2019;29: 2652–2664.e4. doi:10.1016/j.cub.2019.06.078

65. Kirby BB, Takada N, Latimer AJ, Shin J, Carney TJ, Kelsh RN, et al. In vivo time-lapse imaging shows dynamic oligodendrocyte progenitor behavior during zebrafish development. Nat Neurosci. 2006;9: 1506–1511. doi:10.1038/nn1803

66. McGraw HF, Snelson CD, Prendergast A, Suli A, Raible DW. Postembryonic neuronal addition in Zebrafish dorsal root ganglia is regulated by Notch signaling. Neural Dev. 2012;7: 1–13. doi:10.1186/1749-8104-7-23

67. McGraw HF, Nechiporuk A, Raible DW. Zebrafish dorsal root ganglia neural precursor cells adopt a glial fate in the absence of neurogenin1. J Neurosci. 2008;28: 12558–12569. doi:10.1523/JNEUROSCI.2079-08.2008

68. Kimmel CB, Ballard WW, Kimmel SR, Ullmann B, Schilling TF. Stages of embryonic development of the zebrafish. Dev Dyn. 1995;203: 253–310. doi:10.1002/aja.1002030302

